# Newly discovered cichlid fish biodiversity threatened by hybridization with non-native species

**DOI:** 10.1101/2020.08.06.240002

**Authors:** Tabitha Blackwell, Antonia G.P. Ford, Adam G. Ciezarek, Stephanie J. Bradbeer, Carlos A. Gracida Juarez, Benjamin P. Ngatunga, Asilatu H. Shechonge, Rashid Tamatamah, Graham Etherington, Wilfried Haerty, Federica Di Palma, George F. Turner, Martin J. Genner

## Abstract

Invasive freshwater fish systems are known to readily hybridize with indigenous congeneric species, driving loss of unique and irreplaceable genetic resources. Here we reveal that newly discovered (2013-2016) evolutionarily significant populations of Korogwe tilapia (*Oreochromis korogwe*) from southern Tanzania are threatened by hybridization with the larger invasive Nile tilapia (*Oreochromis niloticus*). We use a combination of morphology, microsatellite allele frequencies and whole genome sequences to show that *O. korogwe* from southern lakes (Nambawala, Rutamba and Mitupa) are distinct from geographically-disjunct populations in northern Tanzania (Zigi River and Mlingano Dam). We also provide genetic evidence of *O. korogwe* x *niloticus* hybrids in three lakes and demonstrate heterogeneity in the extent of admixture across the genome. Finally, using the least admixed genomic regions we estimate that the northern and southern *O. korogwe* populations most plausibly diverged approximately 140,000 years ago, suggesting that the geographical separation of the northern and southern groups is not a result of a recent translocation, and instead these populations represent independent evolutionarily significant units. We conclude that these newly-discovered and phenotypically unique cichlid populations are already threatened by hybridization with an invasive species, and propose that these irreplaceable genetic resources would benefit from conservation interventions.

---

Freshwater ecosystems are undergoing rapid changes in biodiversity due to the interacting effects of habitat degradation, over-exploitation, water pollution, flow modification and species invasion (Sala *et al.* 2000; Dudgeon *et al.* 2006; Millennium Ecosystem Assessment, 2016). As human population sizes continue to rise, and climate change becomes an ever-increasing threat, these impacts are predicted to grow (Martinuzzi *et al.* 2014; Arroita *et al.* 2017; Kalacska *et al.* 2017). A specific issue is hybridization between introduced species and native fish species. This has been reported in closely-related species from multiple fish families, including the salmonids (Muhlfield *et al.* 2014; Mandeville *et al.* 2019), cichlids (Firmat *et al.* 2013; Shechonge *et al.* 2018) and cyprinids (Almodóvar *et al.* 2012; Hata *et al.* 2019), and is likely to become increasingly common due to the spread of freshwater species for aquaculture and inland fisheries enhancement (Deines *et al.* 2014). However, the full evolutionary and ecological consequences of hybridization between invasive and native species are typically unclear, and further studies of the impact of hybridization events on native biodiversity are required.

African inland fisheries depend heavily on “Tilapias” (Brummett & Williams, 2000), a group of cichlids that includes the commercially important genera *Oreochromis, Sarotherodon and Coptodon*. Among the most favoured of these species is the Nile tilapia, *Oreochromis niloticus*, which has broad physiological tolerances of environmental conditions, potential for rapid growth, and thus has been widely translocated across the continent (Josupeit, 2010; Dienes *et al.* 2014). However, because of these traits the species is also highly invasive within its introduced range (Ogutu-Ohwayo, 1990; Canonico *et al.* 2005; Deines *et al.* 2017). Moreover, *O. niloticus* is also known to hybridize with native *Oreochromis* species at the locations where it has been introduced, for example with *Oreochromis mossambicus* in Southern Africa (D’Amato, 2007), *Oreochromis esculentus* in Lake Victoria (Angienda *et al.* 2011) and *Oreochromis urolepis* and *Oreochromis jipe* in Tanzania (Shechonge *et al.* 2018; Bradbeer *et al.* 2019). However, despite the growing concern surrounding the impacts of hybridization on native *Oreochromis* populations, the potential loss of unique native genetic diversity due to hybridization with *O. niloticus* remains poorly studied. This is an important area to study because shifts in cichlid fish biodiversity and community composition can lead to fundamental changes in ecosystem functioning (Lévêque 1995), and loss of potential valuable genomic resources for future *Oreochromis* aquaculture strain development (Eknath & Hulata 2009; Lind *et al.* 2012).

Tanzania has a rich diversity of *Oreochromis* species, and preservation of this natural species and its genetic diversity has been recognized as an important conservation goal, given threats of changing environment and hybridization with invasive *Oreochromis* species (Shechonge *et al.* 2018). Recently (between 2013 and 2016) populations of *Oreochromis korogwe* were discovered in three lakes in southern Tanzania near Lindi (Lakes Rutamba, Nambawala and Mitupa; hereafter referred to as ‘southern populations’; Fig. 1). Previously this species was only known from the Pangani and Zigi river catchments in northern Tanzania (hereafter referred to as ‘northern populations’; Fig. 1), some 500 km north of Lindi (Trewavas, 1983; Bradbeer *et al.* 2018; Shechonge *et al.* 2019); the holotype is a specimen from Korogwe in the Pangani catchment (Lowe, 1955). The close evolutionary relationship between representatives of the northern and southern populations has been confirmed in a recent genus-level phylogeny, based on ~3000 bp of nuclear DNA across six loci and ~1500bp of mtDNA (Ford *et al.* 2019, where they were referred to as *O. korogwe* and *O*. sp. Rutamba, respectively). Importantly, the rivers between Lindi and the Pangani are populated naturally only by *O. urolepis*. Such a large geographic discontinuity in the apparent natural distribution of *Oreochromis* is not known in any other species (Trewavas 1983, Shechonge *et al.* 2019), and is rare in other African freshwater fishes (e.g. Skelton 2001). Importantly, in all three of the southern lakes studied, the invasive *O. niloticus* was also found, and the presence of phenotypically intermediate individuals suggested the presence of hybrids.

**Figure 1.**
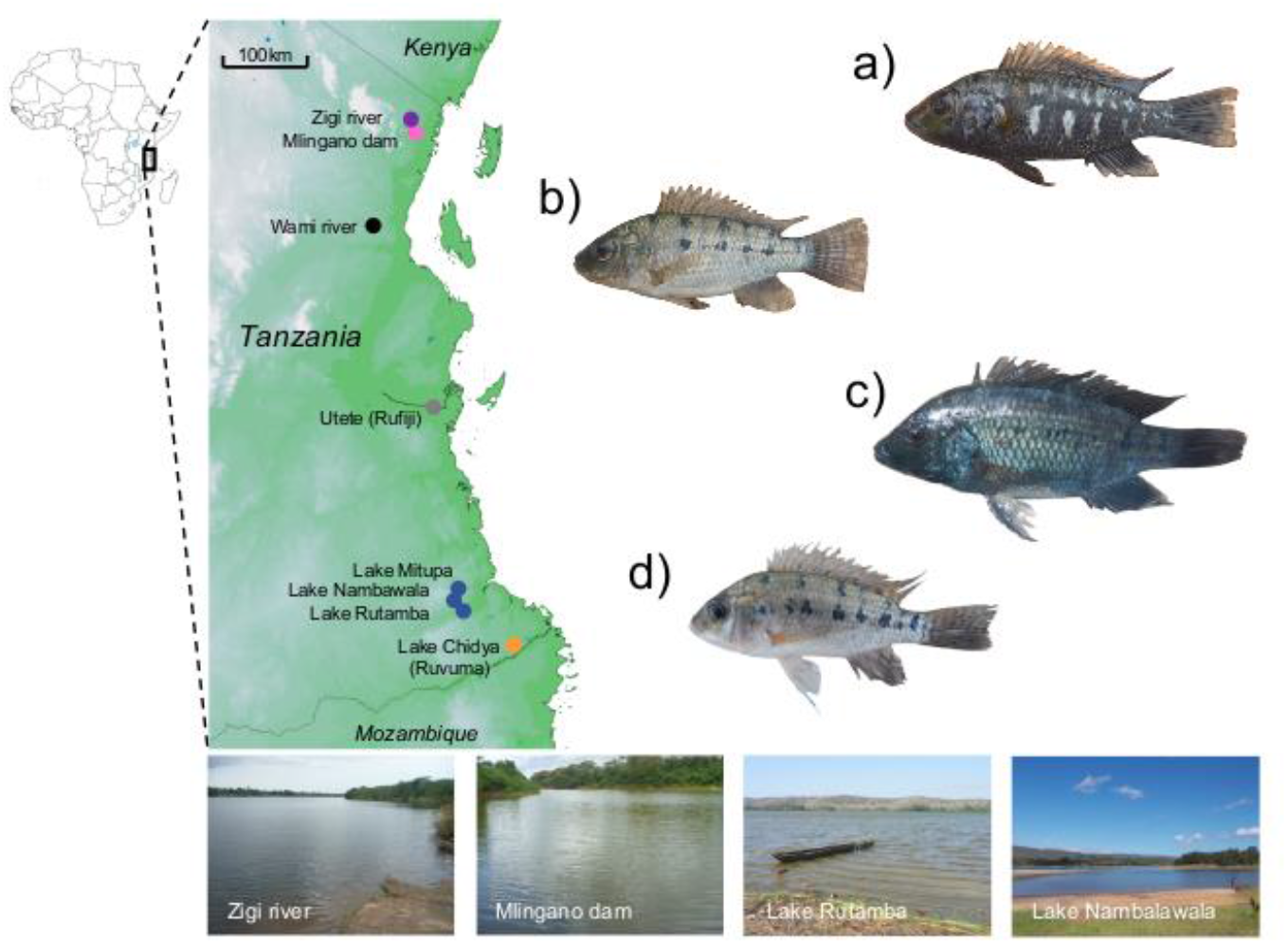
Sampling sites and example specimens of focal populations. a) northern *O. korogwe* male, b) northern *O. korogwe* female, c) southern *O. korogwe* male, d) southern *O. korogwe* female. Pink and purple filled circles indicate northern *O. korogwe* populations sampled, darker blue filled circles locations of the southern *O. korogwe* populations sampled. Grey and black filled circles indicate the sampling locations of *O. urolepis* (Wami and Rufiji and rivers, respectively). The orange filled circles indicate the sampling location of *O. placidus* (Lake Chidya).

In this study we aimed to characterize the diversity and origins of the newly discovered southern populations of *O. korogwe*. We first quantified the extent of hybridization between these populations and invasive Nile tilapia. We then evaluated the possibility that the southern population could be a newly recognized evolutionarily significant unit (*sensu* Fraser & Bernatchez 2001), by comparing genetic and morphological differences with northern *O. korogwe*. We also investigate varying levels of admixture across the genome from *O. niloticus* into southern *O. korogwe*. These results demonstrate that an evolutionarily significant unit is threatened by hybridization with an invasive species, and add to a growing body of evidence for the heterogenous nature of admixture across genomes during hybridization events.

## Materials and Methods

### Study sites and sample collection

*Oreochromis korogwe*, *O. niloticus* and their potential hybrids were collected from southern Tanzania (Lake Rutamba, Lake Nambawala, and Lake Mitupa) on the 14 August 2013, 2-4 May 2015 and 21-27 October 2016 (Fig. 1; Table 1). Samples of *O. korogwe* were collected from northern Tanzania (Zigi River and Mlingano Dam) on the 18 August 2015 (Fig. 1; Table 1). Samples were collected either using multi-mesh gill nets, a seine net, or from purchasing from local fishermen. Multi-mesh nets measured 30m in length with a stretched depth of 1.5m height, and 12 panels each 2.5 meters long. Mesh sizes for panels were in the following order 43mm, 19.5mm, 6.25mm, 10mm, 55mm,Need 8mm, 12.5mm, 24mm, 15.5mm, 5mm, 35mm and 29mm. The seine net measured 30 m in length, 1.5 m in height with 25.4 mm mesh and fine mesh cod end.

**Table 1.**
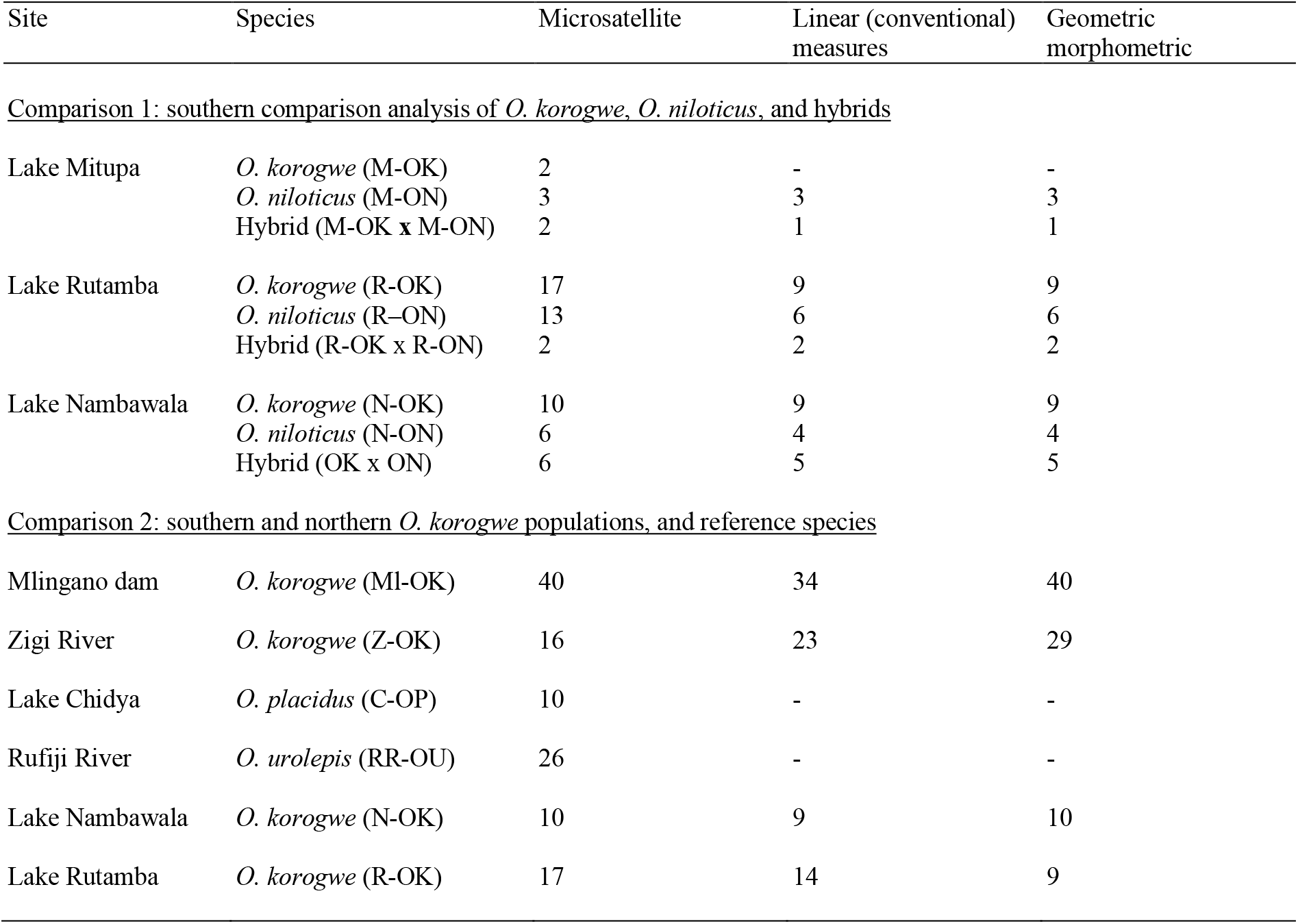
Sample sizes for southern comparison analysis of *O. korogwe*, *O. niloticus*, and individuals of hybrid origin (comparison 1) and comparisons of southern and northern *O. korogwe* populations and reference *O. urolepis* and *O. placidus* (comparison 2).

| Site | Species | Microsatellite | Linear (conventional) measures | Geometric morphometric |
| --- | --- | --- | --- | --- |
| <u>Comparison 1: southern comparison analysis of *O. korogwe*, *O. niloticus*, and hybrids</u> |  |  |  |  |
| Lake Mitupa | *O. korogwe* (M-OK) | 2 | - | - |
|  | *O. niloticus* (M-ON) | 3 | 3 | 3 |
|  | Hybrid (M-OK x M-ON) | 2 | 1 | 1 |
| Lake Rutamba | *O. korogwe* (R-OK) | 17 | 9 | 9 |
|  | *O. niloticus* (R-ON) | 13 | 6 | 6 |
|  | Hybrid (R-OK x R-ON) | 2 | 2 | 2 |
| Lake Nambawala | *O. korogwe* (N-OK) | 10 | 9 | 9 |
|  | *O. niloticus* (N-ON) | 6 | 4 | 4 |
|  | Hybrid (OK x ON) | 6 | 5 | 5 |
| <u>Comparison 2: southern and northern *O. korogwe* populations, and reference species</u> |  |  |  |  |
| Mlingano dam | *O. korogwe* (MI-OK) | 40 | 34 | 40 |
| Zigi River | *O. korogwe* (Z-OK) | 16 | 23 | 29 |
| Lake Chidya | *O. placidus* (C-OP) | 10 | - | - |
| Rufiji River | *O. urolepis* (RR-OU) | 26 | - | - |
| Lake Nambawala | *O. korogwe* (N-OK) | 10 | 9 | 10 |
| Lake Rutamba | *O. korogwe* (R-OK) | 17 | 14 | 9 |

Other samples used for this study were *O. placidus rovumae* from Lake Chidya in the Ruvuma catchment sampled on 18 August 2013, *O. placidus rovumae* from the Ruvuma river sampled on 16 August 2013, *O. placidus rovumae* from the Muhuwesi river (Ruvuma drainage) sampled on 17 August 2013, *O. urolepis* from Lake Lugongwe near Utete on the Rufiji river sampled on 11 March 2015, *O. urolepis* from Mbuyuni pool on the Wami river sampled on 22 August 2015, and *O. niloticus* from within its native (rather than introduced) distribution in Lake Albert, Uganda, sampled on 29 October 2015 (Tables S1, S2). Field collected samples were preserved either in 96-100% ethanol or DMSO salt buffer.

### Population genetics – microsatellite genotyping

DNA was extracted from fin clips using the Wizard kit from Promega (Madison, WI). Samples were genotyped at 13 microsatellite loci (Table S3), sourced from Saju *et al.* (2010) and Liu *et al.* (2013), within two multiplex reactions for each sample. The first contained 6 loci and the second 7 loci. Polymerase Chain Reaction (PCR) was performed using solutions comprising: 1*μ*l DNA, 0.2*μ*l of each 10*μ*M forward primer, 0.2*μ*l of each 10*μ*M reverse primer, 5*μ*l 2x Qiagen Multiplex PCR Master Mix, and made up to 10 *μ*l using RNase-free water. PCR was conducted on a 3PRIME X/02 thermocycler (Techne), with the following settings: an initial denaturation at 95°C for 60 seconds, followed by 35 cycles of 94°C for 30 seconds, 57°C for 90 seconds, and 72°C for 60 seconds. The final extension stage was 60°C for 30 minutes. Products were genotyped on an Applied Biosystems 3500 Genetic Analyser alongside a LIZ500 size standard. Peaks were identified automatically using the software Genemapper v4.1 (Applied Biosystems; CA) and checked manually for accuracy. Arlequin v3.5 (Excoffier and Lischer, 2010) was used to summarize genetic diversity of populations and test for deviations from Hardy Weinberg Equilibrium.

### Population genetics – microsatellite evidence of hybridization in the southern lakes

Potential hybrid individuals between *O. korogwe* and *O. niloticus* were identified from microsatellite data using a two-step process. 1) For all three lakes simultaneously, the find.clusters function in the R package adegenet v2.1.1 (Jombart and Ahmed 2011) was applied, selecting max.n.clust = 40, and the maximum number of principal components, to make a preliminary assignment of individuals to two genetic clusters (*K* = 2), representing *O. korogwe* and *O. niloticus.* 2) Structure v2.3.4 (Pritchard *et al.* 2000) was used to quantify probability of assignment of individuals to the two species. Structure runs used *K* = 2 with the adegenet find.clusters assignments as a prior. The admixture model was used, with each run including 100,000 steps as burn-in, followed by 100,000 sampled steps. Runs were repeated a total of 10 times, and Structure results were summarized across the runs using Clumpak (Kopelman *et al.* 2015), with putatively purebred individuals identified as those possessing > 0.9 probability of belonging to either *O. korogwe* or *O. niloticus*, and the remainder considered to be putative *O. niloticus* x *korogwe* hybrids. To ordinate the genetic structure present within the southern lakes, a Factorial Correspondence Analysis in Genetix v4.05 was used (Belkhir *et al.* 1999).

### Population genetics – microsatellite differences between northern and southern O. korogwe

The genetic structure of putative purebreds from the southern *O. korogwe* populations (Lake Nambawala and Lake Rutamba) to the northern *O. korogwe* populations (Zigi River and Mlingano Dam) was compared, as well as *O. placidus* (Lake Chidya) and *O. urolepis* (Rufiji river at Utete) (Table S4). *Oreochromis korogwe* individuals from Lake Mitupa were not included in the analysis due to the small sample size of purebred individuals (*n* = 6). Structure v2.3.4 (Pritchard *et al.* 2000) was used to assess population genetic structure, using sampling location as a prior. The admixture model was selected, with each run including 100,000 steps as burn-in, followed by 100,000 sampled steps. Runs for each potential number of clusters *K* (between 2 and 6), were repeated a total of 10 times, and the results were summarized using Clumpak (Kopelman *et al.* 2015). Within Clumpak the Evanno method (Evanno *et al.* 2005) was used to identify the optimal number of clusters present in the data. A Factorial Correspondence Analysis in Genetix 4.05 was used to ordinate the genetic structure (Belkhir *et al.* 1999). Genetic structure among the populations was estimated in Genepop v4.2 (Rousset, 2008) using *F*_ST_ and the significance of differences among populations was estimated using Exact tests with default settings.

### Whole genome resequencing - library preparation and data analysis

Twelve samples were processed for whole genome resequencing, comprising two *O. niloticus* specimens, two *O. urolepis* specimens, two *O. placidus* specimens, three specimens from a northern *O. korogwe* population (Mlingano Dam) and three specimens from a southern *O. korogwe* population (Lake Nambawala) (Tables S1 and S2). The selection of these specimens was based on phenotypic characters, and they were all assumed to be purebred at the time of selection for WGS analysis. DNA was extracted from fin clips using a PureLink Genomic DNA extraction kit (ThermoFisher, MA, USA). Genomic libraries were prepared using the Illumina TruSeq HT paired-end read protocol, by Earlham Institute Pipelines department. Samples were sequenced using an Illumina HiSeq 2500 with version 4 chemistry (10 samples per lane; target 5X coverage per sample) and a 125bp paired end read metric. Initial data handling and quality analysis included demultiplexing and conversion to FASTQ files, followed by use of FASTQC (Andrews, 2010) for quality analysis of FASTQ files.

### Whole genome resequencing - Read mapping and SNP calling

Reads were mapped against the “GCF_001858045.2” reference *Oreochromis niloticus* assembly (Conte *et al.* 2019) from NCBI, using the default settings of BWA-MEM v0.7.17 (Li 2013), with the output bam files subjected to samtools v1.9 (Li *et al.* 2009) fixmate prior to being sorted by co-ordinate. Duplicate reads were then marked using picardtools (v1.140; http://broadinstitute.github.io/picard). SNPs were then called using gatk (v4.1.6.0) (McKenna *et al.* 2010). First, HaplotypeCaller was used on each sample, using min-pruning 1, min-dangling-branch-length 1 and heterozygosity 0.01. All samples were collated using GenomicsDBImport, before joint-genotyping with GenotypeGVCFs. SNPs within 5 base pairs of an indel were removed using BCFtools v1.10.2, and then SNPs with total depth exceeding 180 (average exceeding 15x coverage per sample), quality-by-depth less than 2, FS greater than 10, MQ less than 30, MQRankSum less than −2, ReadPosRankSum less than −2 or SOR greater than 3 were filtered using GATK VariantFiltration (Table S5). Individual genotypes with depth less than 3 were replaced with a no-call. BCFtools v1.10.2 was then used to remove sites which overlapped with indels in some samples, and remove SNPs which fell in scaffolds other than the inferred linkage groups.

### Whole genome resequencing – PCA, ADMIXTURE and phylogenetic analysis

For PCA and ADMIXTURE analysis, biallelic SNPs within the linkage groups, with a minor-allele count of at least 3 and less than 25% missing taxa per site were extracted. These were filtered for linkage-disequilibrium using PLINK v2.0.0 (Purcell *et al.* 2007), removing SNPs with r^2^ > 0.2 in sliding windows of 50 SNPs, with 10 SNP overlap. PCA analysis on the resulting 160,883 SNPs was then carried out in PLINK, with the top 20 principal components reported. To investigate population membership, we used Bayesian clustering in ADMIXTURE v1.3.0 (Alexander *et al.* 2009) on the same SNP dataset. which uses a similar algorithm to the Structure program used for the microsatellite analysis, but runs more quickly on large datasets. ADMIXTURE analysis was run using the main algorithm, from *K* = 1 to *K* = 6, with default values for cross-validation error estimation.

For the nuclear phylogeny, SNPs with at least one homozygous reference and one homozygous alternate site were extracted. A phylogenetic tree was inferred using RAxML v8.0.20 (Stamatakis 2014) and the GTRGAMMA model of evolution, with the lewis ascertainment bias correction and 200 rapid bootstraps. To examine the mitochondrial phylogeny, *de novo* assemblies were produced from the raw reads for each individual, using mtArchitect (Lobon *et al.* 2016), which accounts for nuclear mitochondrial DNA segments. These assemblies were aligned using MAFFT v7.271 (Katoh and Standley 2013). A phylogenetic tree was then inferred using RAxML, the GTRGAMMA model of evolution and 200 rapid bootstraps.

### Whole genome resequencing - differentiation across the genome

Relative genetic differentiation between populations (Weir and Cockerham *F*_ST_) as well as absolute sequence divergence within (pi) and between (Dxy) populations were calculated in non-overlapping 50kb windows using popgenWindows.py (https://github.com/simonhmartin/genomics_general). For this analysis, SNPs were filtered to include only sites with at least two individuals per population. Both pi and Dxy require counts of all sites in a window, including SNPs and monomorphic sites. To get the number of callable sites across the genome, we used the CallableLoci function within GATK v3.7.0 (McKenna *et al.* 2010) and a custom script to get counts in each 50kb window. Inferred values of Dxy and pi from popgenWindows.py were then corrected to account for monomorphic sites, which were not in the input vcf, by multiplying them by the number of SNPs in the windows, and then dividing by the total number of callable sites. The *O. placidus* samples were not included as one specimen was evidently a hybrid (see Results).

We also used Twisst (Martin and Van Belleghem 2017) to explore phylogenetic relationships across the genome. Although we did not perform phasing and imputation for the main whole genome dataset analysis due to the small sample size, it is useful for phylogenetic analysis and likely to be accurate over the short (50-SNP) regions considered in the Twisst analysis (discussed further in Martin & Belleghem 2017). We therefore first performed phasing and imputation of biallelic SNPs with minor-allele count of at least three and less than 3 missing taxa using Beagle v4.1 (Browning and Browning 2007) with a window size of 10,000 and overlap of 1000 SNPs. Phylogenetic trees were inferred over sliding 50-SNP windows (requiring at least 40 SNPs per individual), with a 10 SNP overlap using IQtree v1.6.12 (Nguyen *et al.* 2015) using the best fit model for each, with ascertainment bias correction, using scripts modified from genomics_general (https://github.com/simonhmartin/genomics_general).. We then ran Twisst to calculate topology weightings for each window using the method ‘complete’. A smoothing parameter was applied with a loess span of 500,000 base pairs, with a 25,000 spacing.

### Divergence times

We used estimates of Dxy to estimate divergence times between *korogwe* from Mlingano, and Nambawala. To convert estimates of Dxy to divergence times, we used the genome-wide mutation (μ) estimate of 3.5 × 10^−9^ (95% confidence interval: 1.6 × 10^−9^ to 4.6 × 10^−9^) per bp per generation as recently estimated for haplochromine cichlids in Malinsky *et al.* (2018) and assumed a generation time of one year. This was chosen because studies of wild populations of *Oreochromis* species suggest that generation time varies from 3-36 months and is dependent on habitat and population density, with populations in shallow-water and inshore habitats maturing at 12 months or less (Lowe-McConnell 1982). Given the small adult body size of *O. korogwe* and its occurrence in shallow eutrophic water bodies, we used a generation time at the lower end of this range of 1 year.

Estimates of Dxy between the Mlingano and Nambawala *korogwe* will be increased in genomic regions involved with introgression or incomplete lineage sorting. Using the Twisst output, we identified windows where the weighting of the species tree was 1, i.e. there is no evidence for discordance. Using bedtools (v2.28.0) (Quinlan and Hall 2010), we found the 50kb windows overlapping these regions, and used Dxy from these regions to get a measure of divergence in windows supporting the species tree.

### D3 statistics

The genotypes used for sliding window *F*_ST_, Dxy and pi analysis were using to calculate pairwise-distances between each individual, in 50kb non-overlapping windows across the genome, using *distMat.py* (https://github.com/simonhmartin/genomics_general). This pairwise-distance was corrected using the number of callable sites per window (see that section of the methods). *D3* statistics can be used to test for introgression between either P3 and P2 or P3 and P1 in a three-taxon phylogeny (P3,(P2,P1));, without the presence of an outgroup, using genetic distances. Introgression would be expected to result in reduced genetic distance between the two taxon in question. Using the equation *D3*= (*d*P1P3 – *d*P2P3) / (*d*P1P3 + *d*P2P3); where *d*P1P3 is the distance between P1 and P3 and *d*P2P3 is the distance between P2 and P3, a result where *D3* is significantly less than 0 indicates introgression between P1 and P3, whereas a result where *D3* is significantly greater than 0 indicates introgression between P2 and P3 (Hahn and Hibbins 2019). Significance was assessed by 1000 block bootstrap replicates, with the standard deviation used to calculate p values using the overall mean *D3*. The test was carried out between all trios of species where P1 was an individual from *O. korogwe* Nambawala, P2 was an individual from *O. korogwe* Mlingano and P3 was an individual from either *Oreochromis niloticus* or *Oreochromis urolepis*.

### Geometric morphometrics – analyses of individuals from the southern lakes

Ethanol preserved specimens were photographed on their left side in standard orientation with a scale. The image was calibrated for size and 24 landmarks (Fig. S1) were placed onto the image of each specimen using tpsDIG 1.40 (Rohlf, 2004). All microsatellite-genotyped fish (See below) were included in geometric morphometrics, except for specimens of *O. korogwe* where pelvic fins were naturally absent. Landmark data were subjected to a Procrustes analysis in MorphoJ 1.06 (Klingenberg, 2011). Individuals were assigned to one of three groups based on Structure results (purebred *O. niloticus,* purebred *O. korogwe*, hybrid *O. niloticus* x *korogwe*). The Procrustes coordinates were then regressed against centroid size in MorphoJ 1.06, and the size standardized residuals from this regression analysis were then used in a stepwise Discriminant Analysis in SPSS 24 (IBM, London), with purebred *O. niloticus* and purebred *O. korogwe* placed in *a-priori* known categories, and hybrid individuals uncategorized.

Linear morphometric measurements were taken from each genotyped specimen collected in 2016 using digital calipers, following methods outlined in Barel *et al.* (1977) and Snoeks (2004). The following measures were made: standard length, body depth, head length, caudal peduncle length, caudal peduncle depth, dorsal fin base length, anal fin base length, pectoral fin base length, pelvic fin length, caudal fin length, head width, snout length, eye length, interorbital width and lower jaw length. Measurements were log_10_ transformed and size-standardized residuals generated from a linear regression against standard length. Individuals were assigned to the three different groups based on Structure results (purebred *O. niloticus,* purebred *O. korogwe*, hybrid *O. niloticus* x *korogwe*). The size-standardized residuals were used in a Discriminant Analysis in SPSS 24 (IBM, London), with purebred *O. niloticus and* purebred *O. korogwe* placed in *a-priori* known categories, and hybrid individuals remaining uncategorized.

### Morphological comparisons between northern and southern O. korogwe

The morphology of genetically purebred *O. korogwe* from Lakes Rutamba and Nambawala (identified from microsatellite data) was compared to individuals from the Mlingano Dam and Zigi River in northern Tanzania. Geometric morphometric landmarks and linear morphometric measurement data were collected using the methods described above. The geometric morphometric landmarks were subjected to a Procrustes standardization and the resultant Procrustes coordinates were subjected to a pooled within-group regression against centroid size, generating size standardized residuals. These residuals were used in a Canonical Variates Analysis in MorphoJ 1.06, and a Discriminant Analysis in SPSS 24. Linear morphometric measurements were log_10_ transformed. A small number (9 of 2000) of measurements were interpolated using Bayesian PCA in the R package pcaMethods (Stacklies *et al.* 2007), allowing individuals with absent pelvic fins or damaged fins to be included in analyses. We then pooled within-group regressions of each variable against standard length, treating each of the four populations as a group. The size-standardized residuals generated from these regressions were then used in a Discriminant Analysis in SPSS 24.

## Results

### Population genetics - microsatellite analysis of purebred and hybrid Oreochromis in southern lakes

Using Structure, the majority of individuals were assigned to one of the two parent species with a probability of >90%. Individuals that were not able to be assigned to a single species with a probability of >90% were considered hybrids. In total these hybrids comprised 29% of individuals sampled from Lake Mitupa (2 of 7), 27% of individuals from Lake Nambawala (6 of 22), and 6% of individuals from Lake Rutamba (2 of 32) (Fig. 2a,b).

**Figure 2.**
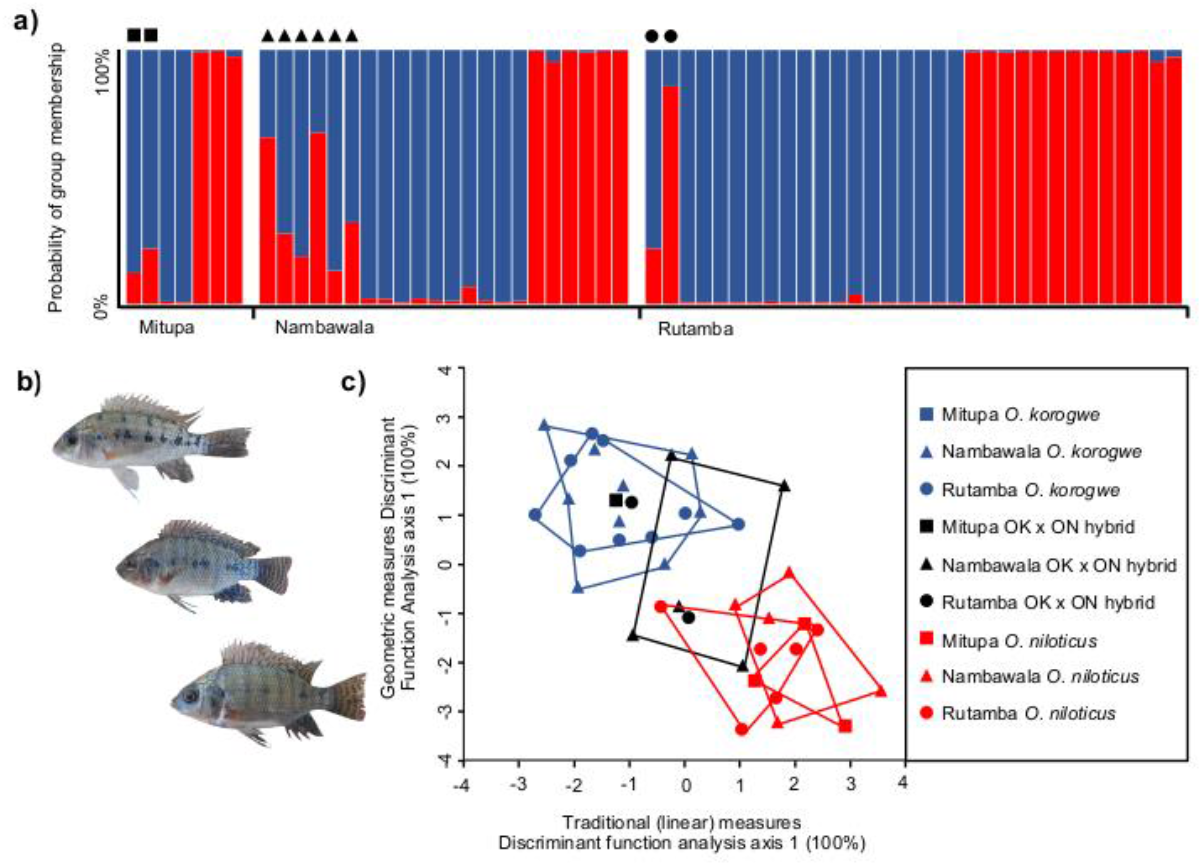
Genetic and morphological contrasts between *O. korogwe, O. niloticus* and *O. korogwe* x *niloticus* hybrids. a) Structure assignment of individuals to populations (*K* = 2) using microsatellite data from *Oreochromis* from the southern lakes. Filled black symbols indicate individuals of putative hybrid origin. b) images of *O. korogwe* (top), *O. korogwe x niloticus* (middle) and *O. niloticus* (bottom). c) Discriminant function axes illustrate distinctive morphology of purebred *O. korogwe* (blue)s and *O. niloticus* (red) *O. korogwe x niloticus* hybrid individuals which overlap in morphospace with parent taxa.

### Morphological comparisons of purebred and hybrid Oreochromis in southern lakes

Discriminant Analysis of geometric morphometric data demonstrated that *O. niloticus* and *O. korogwe* individuals could be reliably separated (Wilk’s λ = 0.272, χ^2^ = 37.054, *P* < 0.001) with 30 of 32 purebred individuals correctly classified (Table S6). Equally, Discriminant Analysis using linear morphometric measurements showed that that *O. niloticus* and *O. korogwe* individuals could be reliably separated (Wilk’s λ = 0.314, χ^2^ = 32.401, *P* < 0.001), with 29 of 32 purebred individuals correctly classified (Table S6). Typically, *O. niloticus* were characterized as possessing a longer and broader head (Table S7). Hybrid morphospace overlapped with that of purebred species in both datasets (Fig. 2c).

### Population genetics – microsatellite genetic structure among Oreochromis populations

Structure analyses indicated the optimum number of genetically distinct populations across the six sampled populations was *K* = 5, with the southern populations from neighbouring lakes Rutamba and Nambawala resolved as genetically homogeneous group (Fig 3a). All *O. korogwe* were genetically distinct from reference populations of *O. urolepis* from the Rufiji river and *O. placidus* from Lake Chidya in ordination plots (Fig. 3b). Analysis including only *O. korogwe* revealed the Zigi river and Mlingano dam populations to be distinct from one another, and to both populations from the south (Fig. 3c). In pairwise comparisons, all populations showed highly significant genetic differences, with exception of *O. korogwe* from Lakes Rutamba and Nambawala (Table 2). No populations showed clear patterns of significant deviation from Hardy-Weinberg Equilibrium in microsatellite loci (Table S4).

**Table 2.** Matrix of *F*_ST_ pairwise comparisons (below left) and corresponding *P* values from Exact tests (above right).

|  | *O. placidus*<br>Chidya | *O. korogwe*<br>Zigi | *O. korogwe*<br>Mlingano | *O. urolepis*<br>Rufiji | *O. korogwe*<br>Rutamba | *O. korogwe*<br>Nambawala |
| --- | --- | --- | --- | --- | --- | --- |
| *O. placidus* Lake Chidya |  | <0.001 | <0.001 | <0.001 | <0.001 | <0.001 |
| *O. korogwe* Zigi river | 0.547 |  | <0.001 | <0.001 | <0.001 | <0.001 |
| *O. korogwe* Mlingano dam | 0.761 | 0.341 |  | <0.001 | <0.001 | <0.001 |
| *O. urolepis* Rufiji river | 0.229 | 0.455 | 0.612 |  | <0.001 | <0.001 |
| *O. korogwe* Lake Rutamba | 0.659 | 0.358 | 0.378 | 0.511 |  | 0.473 |
| *O. korogwe* Lake Nambawala | 0.618 | 0.415 | 0.470 | 0.461 | 0.011 |  |

**Figure 3.**
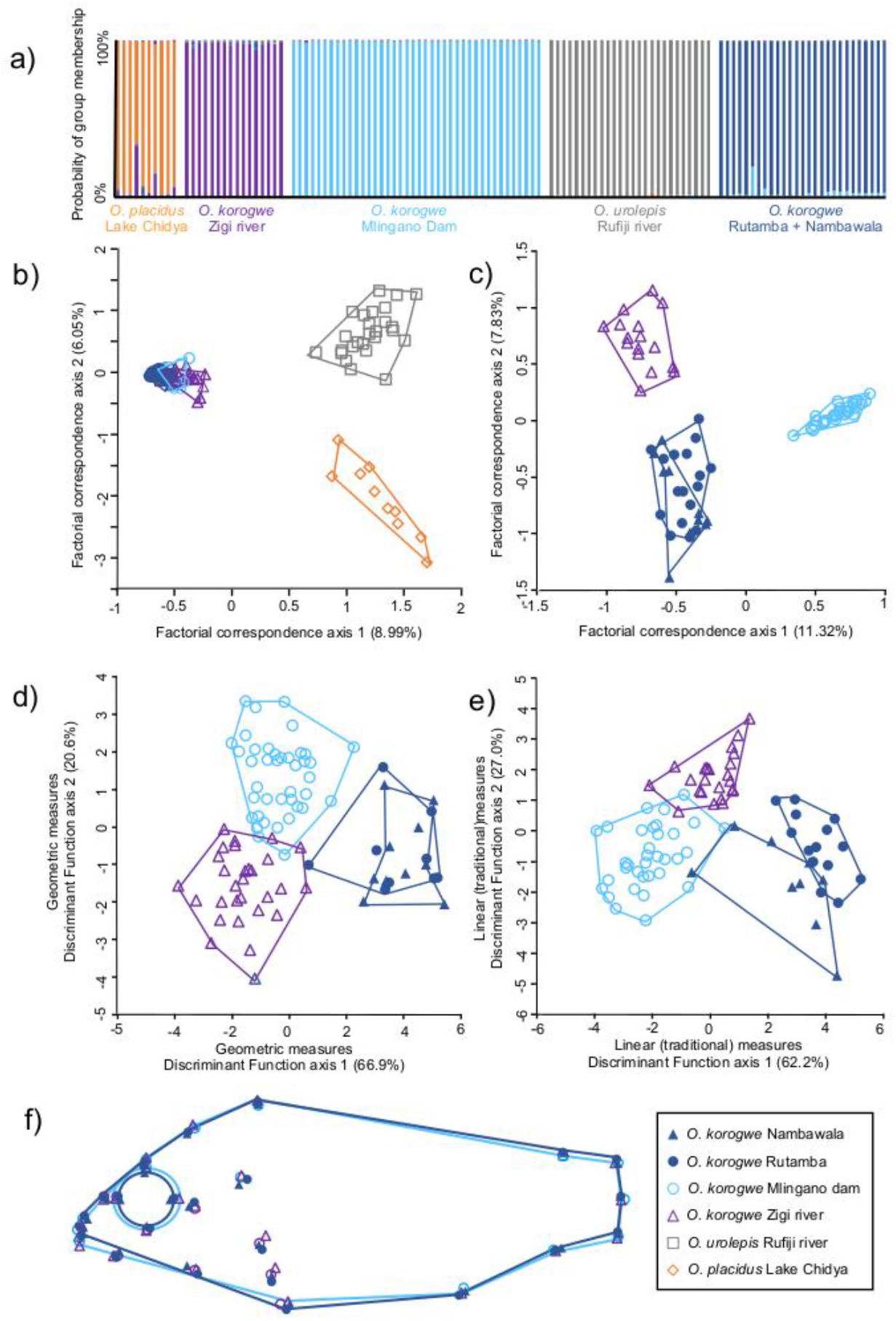
Genetic and morphological analysis of focal populations of *O. korogwe*, and reference populations of *O. urolepis* (Utete), and *O. placidus* (Lake Chidya). a) Structure analysis of the six populations, using *K* = 5. b) Factorial correspondence analysis (FCA) of all populations from all six sites, c) FCA of the four *O. korogwe* populations, d-e) Discriminant Function analysis (DFA) of the four *O. korogwe* populations using linear and geometric measures respectively, and f) Wireframe analysis from Canonical Variates Analysis (CVA) showing geometric morphometric divergence between northern (light blue lines) and southern (dark blue lines) populations.

### Morphological comparisons of northern and southern O. korogwe

Discriminant Function Analysis of both the geometric morphometric data and the traditional morphometric data demonstrated highly significant differences between the northern and southern *O. korogwe* groups (Fig. 3d,e), with the majority of individuals being able to be classified by sampling site using either linear traditional measurement data (74 of 80 individuals), or geometric morphometric data (84 of 88 individuals; Table S8). Discriminant Function Axis 1 separated northern and southern populations in both morphological datasets. In the linear measurements this axis indicated *O. korogwe* from the northern populations to have shallower body depth, a less deep caudal peduncle, a narrower interorbital width and shorter pectoral fins, relative to southern populations (Table S9). Wireframe diagrams indicated northern *O. korogwe* populations had smaller eyes and shallower body dimensions than southern populations (Fig. 3f).

### Whole genome resequencing: phylogenomic analyses

Illumina sequencing resulted in an average of 22 million reads per sample (range: 20.53 to 24.40 million), and mapping rates to the *O. niloticus* reference genome of 97.39 to 99.18% (Table S1). Mean sequencing coverage across the dataset was 5.29X, with approximately half the genome covered with a sequencing depth of at least 5X (Table S1). The filtered datasets and number of SNPs used for downstream analysis are given in Table S10. ADMIXTURE analysis of all 12 samples suggested cross-validation minima at *K* = 2 and *K* = 5, indicating the most likely number of clusters in the dataset (Fig. S2). At *K* = 5, there was a clear separation of the northern and southern *O. korogwe* populations alongside the other species, supported by groupings in PCA (Fig. 4a). The ADMIXTURE analysis also indicated that one *O. placidus* sample was likely an early-generation *O. placidus x niloticus* hybrid or backcross, with approximately 40% *O. niloticus* cluster membership (Fig. 4b).

**Figure 4.**
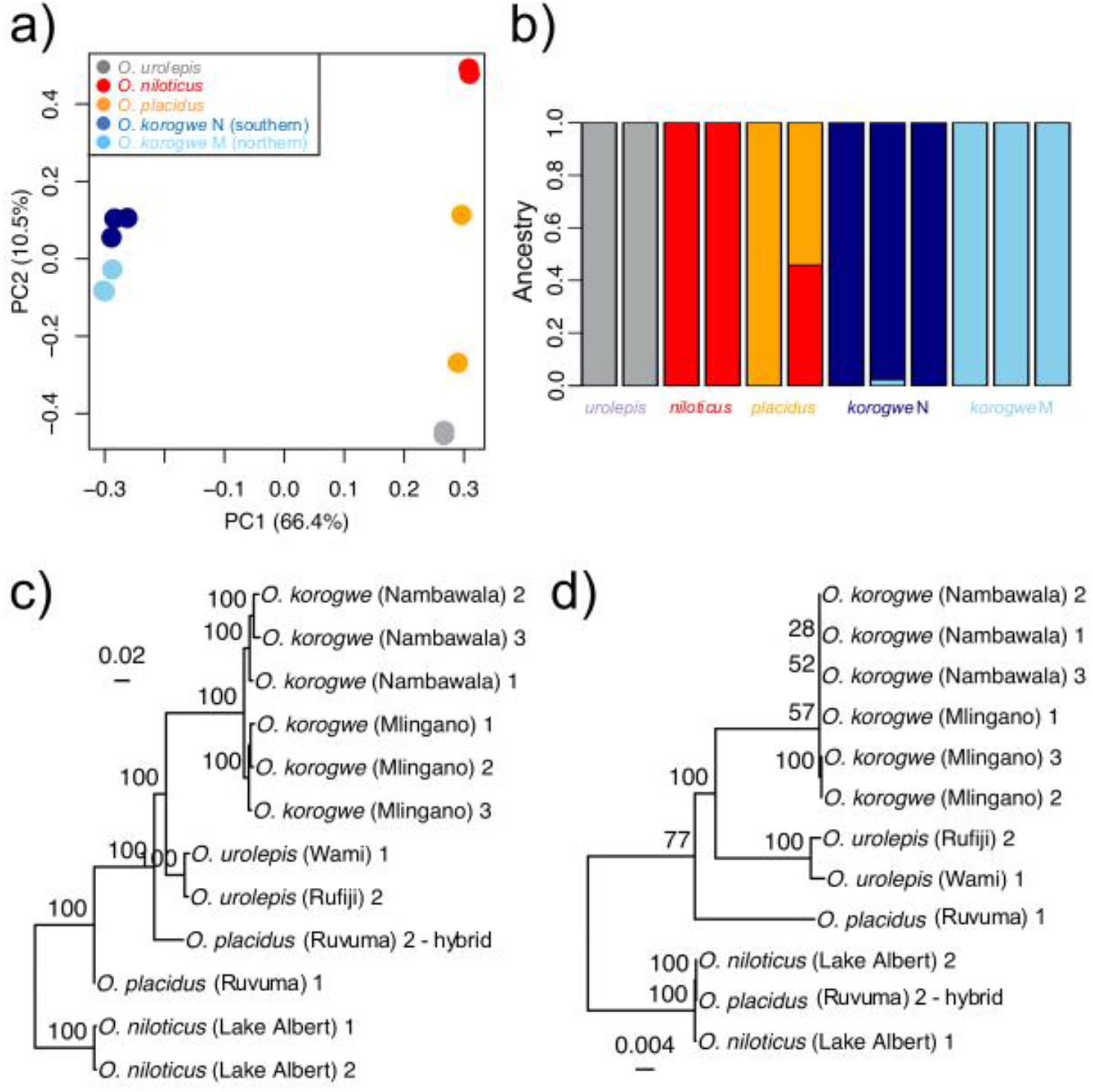
Analyses of genome-wide data. a) Principal Component Analysis (PCA) of all variants, b) Admixture analysis of all variances, c) phylogeny based on nuclear genome variants, using RAxML GTR+ Γ model. d) phylogeny based on mitochondrial genome variants, using RAxML GTR+ Γ model. Scale bars in changes per bp. Values on nodes indicate bootstrap support values for 1000 bootstraps, those >70% shown.

Maximum likelihood phylogenetic analysis indicated that the *O. placidus* hybrid was likely the result of a female *O. niloticus* x male *O. placidus* cross, as the (maternally inherited) mtDNA of the sample clustered with *O. niloticus* (Fig. 4d). Otherwise, there was a clear separation of *O. urolepis*, *O. niloticus*, *O. placidus* and the two *O. korogwe* populations in both the nuclear and mtDNA phylogenies (Fig. 4c-d).

Differentiation (*F*_ST_) was highest among interspecific comparisons (Fig. 5a-f). Between the northern (Mlingano Dam) and southern (Nambawala) *O. korogwe* populations, most 50kb windows had low differentiation, but there were prominent regions of the genome showing very high *F*_ST_ differentiation (Fig. 5e). Notably, there were regions of relatively low genetic differentiation between the *O. niloticus* and *O. korogwe* sampled from Nambawala where the two species are sympatric (Fig 5f), but these were not apparent in the comparison between the fully allopatric *O. niloticus* and *O. korogwe* from Mlingano Dam (Fig. 5d). Section of low *F*_ST_ were also apparent in the comparison of *O. korogwe* from Nambawala and *O. urolepis*. Sections of low *F*_ST_ were showed no clear pattern of being associated with areas of elevated or depleted genomic diversity (pi) in the focal species (Fig. S3). However, it was notable that in all species LG3 had substantially higher variability in genetic diversity relative to other linkage groups, and possessed higher absolute sequence divergence in both intraspecific and interspecific comparisons (Fig. S4).

**Figure 5.**
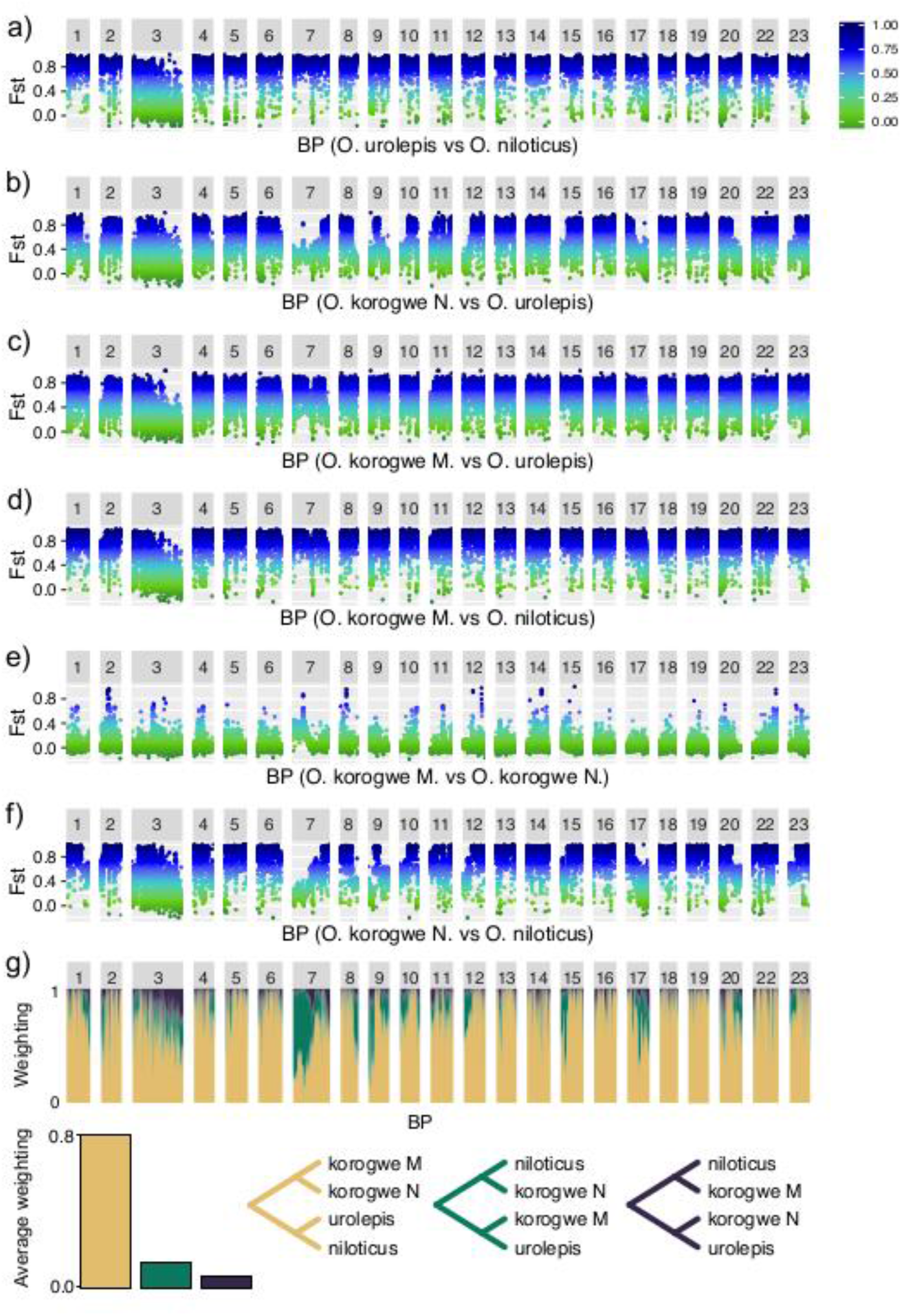
a-f) Pairwise sliding window *F*_ST_ between populations across genome linkage groups, in 50-kb windows, between combinations of *O. niloticus*, *O. urolepis*, southern *O. korogwe* N (Lake Nambawala), northern *O. korogwe* M (Mlingano Dam). g) Phylogenetic representation across genomes of four populations, as estimated by Twisst. Three possible phylogenies for the four taxa are illustrated below, and their colours correspond to relative weightings in plot above. The linkage groups are labelled according to the numbering of the linkage groups in the reference genome.

### Whole genome resequencing: differentiation across the genome and timescale of divergence

Phylogenetic relationships across the genome, generated using Twisst, provided evidence of admixture that was heterogeneous across the genome (Fig. 5g). The two *O. korogwe* populations were resolved as sister taxa across most of the genome. However, for substantive sections of the genome, a phylogeny supported *O. niloticus* and the southern *O. korogwe* (Nambawala) as sister taxa, and *O. urolepis* and northern *O. korogwe* (Mlingano Dam) as sisters. Notably, these tracts of the genome consistent that are consistent with interspecific hybridization corresponded with both the low *F*_ST_ regions *O. niloticus* and the southern *O. korogwe* (Nambawala) (Fig 5f), and low *F*_ST_ region between *O. urolepis* and the northern *O. korogwe* (Mlingano) (Fig 5b). *D3* statistics consistently provided strong statistical support for scenarios of both decreased genetic distance between *O. niloticus* and southern *O. korogwe* in Nambawala compared to between *O. niloticus* and northern *O. korogwe*, and between *O. urolepis* and the northern *O. korogwe* at the Mlingano Dam compared to between *O. urolepis* and southern *O. korogwe* (Table S11).

Overall absolute sequence divergence (Dxy) between the northern (Mlingano Dam) and southern (Nambawala) *O. korogwe* populations was 0.0009 (Fig. S5). Applying the genome-wide mutation (μ) rate estimate of 3.5 × 10^−9^ (95% confidence interval: 1.6 × 10^−9^ to 4.6 × 10^−9^) from Malinsky *et al.* (2018), with a generation time of one year, gave a genome-wide divergence time estimate of 271 KYA (95% CI: 206-594 KYA). Using only those regions of the genome consistent with the hypothesis of the northern and southern *O. korogwe* being sister taxa, the overall absolute sequence divergence (Dxy) was 0.0005, providing a divergence time estimate of 144 KYA (95% CI: 109-315 KYA).

## Discussion

### Population structure of southern and northern O. korogwe

This study confirmed the distinctness of all sampled *O. korogwe* populations from two other species of *Oreochromis* naturally present in coastal rivers of Tanzania, namely *O. placidus* and *O. urolepis*. The results also demonstrated a close evolutionary relationship between *O. korogwe* individuals in northern and southern Tanzania. Nevertheless, there has been extensive morphological divergence between the northern and southern *O. korogwe*, and based on least admixed sections of the genome, this divergence took place approximately 140,000 years ago. Therefore, the data are consistent with these taxa representing independent evolutionarily significant units. The presence of a 500 km gap between the sampled northern and southern populations of *O. korogwe* in Tanzania, is intriguing. In tilapiine cichlids the presence of such gaps is typically due to human intervention. For example, stocking has resulted in *O. niloticus* having a broad discontinuous distribution across Africa, and further afield (Deines *et al.* 2014). However, our results are consistent with the current distribution of *O. korogwe* being natural. The distribution may have arisen from a natural long-distance colonization event, or perhaps that the species once had a wider distribution that has been disrupted through either extirpation or introgression with *O. urolepis*, a species that neatly fits the gap between northern and southern *O. korogwe* (Ford *et al.* 2019; Shechonge *et al.* 2019).

### Morphological variation among O. korogwe populations

Our results showed that the northern and southern *O. korogwe* populations are largely distinct in characters such as body depth, fin length and eye size morphology. The populations are sufficiently divergent in morphology to warrant consideration of these as distinct species under morphological species concepts. The anatomical divergence may be accompanied by ecological differences, as variation in craniofacial morphology and body shape are often related to resource use patterns in cichlids. For example, variation in eye size is related to visual environment (Hahn *et al.* 2017), and fin morphology is related to patterns of habitat use (Colombo *et al.* 2016). Little is known about the feeding habits of *O. korogwe* and detailed analysis of diets and foraging environments within the sampled locations are required to explore functions of the morphological variation observed. Given the allopatric nature of the populations, further ecologically and developmentally-focussed work would also help to reveal if the observed divergence can be attributed to fixed genetic differences, or alternatively variation between environments during development (Parsons *et al.* 2011; Schneider and Meyer, 2017).

Our microsatellite-based results also confirmed the presence of hybrids between *O. korogwe* and invasive *O. niloticus* in all three of the southern lakes, with a frequency of between 6 and 29% of sampled individuals. This level of hybridization is likely to be an underestimate if purebreds are present (Boecklen & Howard, 1997), which our genome-wide analyses also support. Such hybridization between native and non-native species commonly occurs when invader is closely-related to the native species, and the species pair are still reproductively compatible due to an absence of strong reproductive barriers that typically isolate naturally sympatric taxa (Horreo *et al.* 2011, Gainsford, 2014). It is not fully understood what factors influence the extent of reproductive isolation among *Oreochromis* species. However, it is notable that like many African mouthbrooding cichlids, *Oreochromis* exhibit traits indicative of sexual selection based on male colours or the characteristics of breeding territory (Trewavas 1983). It is possible that in this case hybridization between *O. korogwe* or *O. niloticus* takes place due to both species possessing dark male breeding colours (Genner *et al.* 2018). Female mating decisions also biased towards larger individuals in *Oreochromis* species, most likely due to the influence of male-male competition on breeding territory acquisition (Nelson 1995; Fessehaye *et al.* 2006). Hence, is also conceivable that larger *O. niloticus* males have effectively excluded smaller *O. korogwe* males from suitable breeding habitats; but detailed survey and experimental work is required to test this hypothesis, including tests of sex-biases in the direction of hybridization (e.g. Hayden *et al.* 2010; Rognon & Guyomard, 2003).

### Heterogeneity of admixture across the genome

We conducted genome-wide scans of *F*_ST_ and Dxy between *O. niloticus*, *O. urolepis* and *O. korogwe* populations. *F*_ST_ between the northern (Mlingano Dam) and southern (Nambawala) *O. korogwe* populations was typically low across all linkage groups, with peaks of high *F*_ST_ that may reflect genomic regions under directional selection. These peaks of the *F*_ST_ were not clustered, and these regions associated loci associated with the divergent phenotypes of these populations. These patterns are characteristic of early stage speciation under geographical isolation (Seehausen *et al.* 2014).

Between *O. urolepis* and *O. niloticus* a consistent pattern of high *F*_ST_ was present, reflecting the long divergence. On linkage group 3, *F*_ST_ was lower, and but it is notable that this shows an unusually high level of sequence diversity in all our studied *Oreochromis* populations (Fig. S4), as well as a high level of absolute sequence divergence between all populations (Fig. S5). On account of this linkage group being 2-3 times larger than any other in the *Oreochromis* genome (Fig. 6; Conte *et al.* 2019), LG3 has been referred to as a megachromosome, and is likely to consist of a fusion with an ancestral B-chromosome (Conte *et al.* 2020). It is rich in long-coding RNA, genes related to immune response and regulation, and repetitive elements. It has also been reported as containing a sex-determination locus in *Oreochromis*, albeit not in *O. niloticus* itself (Conte *et al.* 2020). Collectively, the high genetic diversity of this linkage group explains the relatively low *F*_ST_ observed between *O. urolepis* and *O. niloticus*, and between other species pairs.

In comparisons between *O. niloticus* and southern *O. korogwe* from Lake Nambawala, there was considerable heterogeneity in *F*_ST_ across the genome. There were notable long-tracts of relatively low *F*_ST_, most conspicuously on linkage groups 1, 7, 9 10, 17, 20 and 23. Many of these were paralleled by low *F*_ST_ between *O. urolepis* and *O. korogwe* from Lake Nambawala. However, the regions of low differentiation were not present in comparisons between *O. niloticus* and northern *O. korogwe* from the Mlingano Dam, or between *O. urolepis* and *O. korogwe* from the Mlingano Dam. This is suggestive of the observed patterns of substantive genomic heterogeneity being reflective of admixture events in the south of Tanzania, after the split from northern *O. korogwe* approximately 140,000 years ago.

Given our microsatellite evidence of individuals of *O. korogwe x niloticus* hybrid ancestry within Lake Nambawala, tracts of low *F*_ST_ between *O. korogwe x O. niloticus* plausibly reflect hybridization between in the southern region. The analysis of phylogenetic relationships of the focal populations in this study using Twisst show that although the species tree relationship is most common across the genome, there is a substantial difference in the frequency of the two discordant relationships, which under incomplete lineage sorting alone would be expected to have the same frequency, The observed excess of the discordant topology grouping *O. niloticus* with *O. korogwe* Nambawala and *O. urolepis* with *O. korogwe* Mlingano (green in Figure 5g) therefore suggests introgression between *O. niloticus* and *O. korogwe* Nambawala or between *O. urolepis* and *O. korogwe* Mlingano. Supporting this, all *D3* analysis suggest significantly lower genetic distances between *O. niloticus* and *O. korogwe* Nambawala and between *O. urolepis* and *O. korogwe* Mligano, than otherwise expected under a model of no-hybridization. However, this three-taxon analysis can be confounded by introgression events involving taxa that have not been included in the analysis. Introgression between *O. niloticus* and *O. korogwe* Nambawala, for example, would increase average the genetic distance between *O. korogwe* Nambawala and *O. urolepis*, as the genetic distance between *O. urolepis* and *O. niloticus* is greater than between *O. urolepis* and *O. korogwe* Nambawala. A single introgression event, between *O. niloticus* and *O. korogwe* Nambawala, could therefore explain both positive results.

The genomic regions of this introgression highlighted by the Twisst analysis overlap with the low *F*_ST_ regions between *O. niloticus* and *O. korogwe* Nambawala, but such low *F*_ST_ regions are not observed between *O. urolepis* and *O. korogwe* Mlingano. The most congruent interpretation of these *F*_ST_ results is introgression between *O. niloticus* and *O. korogwe* Nambawala. The parallel regions of low *F*_ST_ present between *O. korogwe* from Lake Nambawala and *O. urolepis* are unusual however, given that *O. urolepis* has never been recorded inside Lake Nambawala, or elsewhere in the known range of *O. korogwe* (Shechonge *et al.* 2019). One possible explanation for this pattern is that the introduced *O. niloticus* population in Lake Nambawala could itself comprise *O. urolepis* x *niloticus* hybrids, as these species are known to hybridise elsewhere in Tanzania (Shechonge *et al.* 2018), and it is plausible that Nambawala was stocked from a hybrid population. Alternatively, these low *F*_ST_ tracts may reflect recent admixture of ancestral variation shared by both *O. urolepis* and *O. niloticus*. We have not sequenced the *O. niloticus* from Lake Nambawala to test for the presence of recent introgression with *O. urolepis*, but this may be enlightening. We must also note that the low sample sizes (n=2 to 3 individuals) will have limited the accuracy and reliability of *F*_ST_, Dxy and pi statistics. Further studies with more comprehensive phylogenetic and population sampling with greater sample sizes may be able to untangle the nature of introgression events with more precision.

Extensive heterogeneity in the extent of admixture across genomes has been reported in multiple studies of closely related species, including trees (Wang *et al.* 2020), insects (Martin *et al.* 2019, Valencia-Motoya *et al.* 2020) and cichlid fish (Gante *et al.* 2016, Svardel *et al.* 2020). Tracts of the southern *O. korogwe* genome with extensive evidence for hybridization (e.g. LG7, LG9 and LG17), may have resulted from introgressed alleles in those regions being favoured by selection. In North America hybridization between introduced rainbow trout (*Oncorhynchus mykiss*) and native westslope cutthroat trout (*Oncorhynchus clarkii lewisi*), has led to multiple genomic variants being shared between the species, with selection repeatedly favouring some introduced alleles within the native species (Bay *et al.* 2019). Adaptive introgression has similarly been suggested to have led to multiple beneficial traits arising from close-relatives in many species groups, including Darwin’s finches (Lamichhaney *et al.* 2015), snowshoe hares (Jones *et al.* 2018) and multiple plant taxa (Suarez-Gonzalez *et al.* 2018).

In comparisons of *O. korogwe* from Lake Nambawala and *O. niloticus*, regions of the genome with low levels of introgression (e.g. LG6, LG16 and LG19). This may be due to the presence of “barrier” loci that reduce gene flow and maintain species boundaries (Elmer *et al.* 2019). It is been shown that hybridization can suppress recombination rates in some genomic regions of hybrid trout (Ostberg *et al.* 2013). It has also been proposed that recombination is particularly strongly suppressed near genes associated with reproductive isolation among parent species, due to hybrids have a low relative fitness (Hvala *et al.* 2018). In particularly, hybridization could lead to the breakup of coadapted “supergene” clusters, leading to low fitness hybrids, and so these large genomic regions would in principle be among most resistant to introgression. Positive associations between recombination rate of genome and admixture have been described in humans and swordtail fishes (Schumer *et al.* 2018), as well as sympatric pairs of *Heliconius* butterflies (Martin *et al.* 2019). However, accurate estimations of recombination rate require genotype data from more extensive population sampling than has been undertaken for our study, so this remains an untested yet plausible explanation for at some of the heterogeneity observed.

### Conservation implications

Our results support the concept that the northern and southern *O. korogwe* populations are long-diverged and phenotypically-divergent evolutionarily significant units. These may require consideration as discrete species, which will have implications for the biodiversity of tilapias of East Africa. However, the results also illustrate that genetic structure within the newly discovered populations of *O. korogwe* has already been impacted by the invasive species *O. niloticus*. Similarly, the results also show *O. niloticus* has hybridized with *O. placidus* in the neighbouring Ruvuma drainage. Species introductions can have non-reversible impacts on genetic diversity (Dudgeon *et al.* 2006), and therefore the presence of this highly invasive species in these lakes is of considerable concern for the long-term conservation status for these populations. Hybridization could have larger impacts on the genetic diversity of this population over time, especially given evidence from other lakes where *O. niloticus* have been introduced (e.g. Deines *et al.* 2014) and given the lack of understanding of the long-term fitness consequences of these interaction. Although there is some evidence that hybridization could introduce advantageous alleles into the population, our findings suggest that these southern *O. korogwe* populations are likely to be locally adapted to the southern lakes. Therefore, introgression may have negative outcomes for the genetic uniqueness of the *O. korogwe* populations at least.

Our results clearly demonstrate an ongoing threat to unique southern *O. korogwe* populations, and long-term monitoring of the genetic and phenotypic diversity within the studied lakes will yield insights into changes of their status. We suggest that clear conservation actions could be implemented. Given the removal of *O. niloticus* from the southern lakes would be impractical, conservation of the unique genetic resources within the southern lakes would be best done through the identification of potential ark sites. For this research we sampled three of the water bodies in close proximity to the towns of Lindi and Rutamba, and it is possible that *O. korogwe* populations unaffected by *O. niloticus* are present in four additional proximate water bodies that we have not yet been surveyed. Each of these potential ark lakes will need to be intensively investigated to determine the species of fish present, and the potential for *O. niloticus* colonisation via natural waterways. In the absence of the suitable ark sites, the *ex-situ* conservation could be implemented. In both conservation strategies, genome-wide sequencing would be useful to confirm the genetic purity of the stocks, as this study has shown a clear signal of introgression in individuals of *O. korogwe* from Lake Nambawala that were assumed to purebred on the basis of the phenotypes. Therefore, this study underlines the value of using genome-wide sequencing for assessing the conservation status of taxa under threat from hybridization with introduced species.

## Acknowledgements

The work was funded by Royal Society-Leverhulme Trust Africa Awards AA100023 and AA130107 to MJG, BPN and GFT, BBSRC award BB/M026736/1 to GFT, MJG and FdP, and and BBSRC award BB/P016774/1 to WH and FdP We thank the Tanzania Commission for Research and Technology (COSTECH) for fieldwork approval and permits, and staff of the Tanzania Fisheries Research Institute for contributions to fieldwork. We thank Nasser Kazosi for help with sample collection in Uganda, made under permit number IMP/GEN/2014/06. This research was supported in part by the NBI Computing infrastructure for Science (CiS) group through use of the CiS high-performance computing cluster for the analysis of the whole genome resequence data.

## Author Contributions

GFT, MJG and FDP conceived the study. MJG, GFT and BPN designed fieldwork and sampling. TB, SJB, AGPF, CAGJ, BPN, AS, GFT, RT and MJG conducted or supervised fieldwork, or collected data. TB, AGPF, AGC, MJG, GE and WH designed and performed the analysis. TB, AGPF and MJG wrote the first draft of the manuscript. All authors commented on and edited the final manuscript.

## Data Accessibility Statement

- Microsatellite genotype data - DOI: 10.5523/bris.2ka0x87ea99pk221rw3dahmzbn
- Morphological data - DOI: 10.5523/bris.2ka0x87ea99pk221rw3dahmzbn
- DNA resequencing data (raw reads) - to be deposited at the European Nucleotide Archive; Project number: PRJEB36772 on acceptance
- DNA resequencing data (vcf files) – to be deposited at the University of Bristol RDSF on acceptance.

