## Supplementary Material for "Newly discovered cichlid fish biodiversity threatened by hybridization with non-native species"

**for:**

**Table of Contents**

| **Supporting Text.** Commands for whole genome resequence (WGR) data analysis | 2 |
| --- | --- |
| **Table S1**: Whole genome resequencing sample details | 4 |
| **Table S2**: Samples used in each of the analyses | 5 |
| **Table S3**: Microsatellite loci primer sequences | 11 |
| **Table S4**. Genetic (microsatellite) diversity of the focal populations | 12 |
| **Table S5**. WGR quality filtering thresholds applied to the SNP and indel datasets. | 13 |
| **Table S6**. Classification results from Discriminant Function Analysis I | 14 |
| **Table S7**. Correlation of traits with Discriminant Function Axis I (Analysis I) | 14 |
| **Table S8**. Classification results from Discriminant Function Analysis II | 15 |
| **Table S9.** Correlation of traits with Discriminant Function Axes (Analysis II) | 15 |
| **Table S10**. Data files number of SNPs by analysis. | 16 |
| **Table S11**. Results of the D3 statistics | 17 |
| **Figure S1.** Landmarks used in the geometric morphometric analysis. | 18 |
| **Figure S2.** WGR population genetic and phylogenetic analysis. | 19 |
| **Figure S3.** Within population nucleotide diversity (pi) across linkage groups | 20 |
| **Figure S4**. Absolute genetic divergence (Dxy) between population pairs | 21 |

### Supporting Text. Commands for whole genome resequence (WGR) data analysis

Genotyping (Per sample HaplotypeCaller):

gatk --java-options "-Xmx30g" HaplotypeCaller -R $ReferenceO -I $inbam -O 1_HaploCaller/$outname -ERC GVCF --min-pruning 1 --min-dangling-branch-length 1 --heterozygosity 0.01 -G StandardAnnotation -G AS_StandardAnnotation --native-pair-hmm-threads 4

Joint Genotyping:

gatk --java-options "-Xmx100g" GenomicsDBImport -R $Reference --genomicsdb-workspace-path GenDB --intervals intervals.list --max-num-intervals-to-import-in-parallel 30 --overwrite-existing-genomicsdb-workspace --tmp-dir gatktmp $(ls 1_HaploCaller/*.vcf.gz | sed 's/^/-V /g' | tr '\n' ' ')

gatk --java-options "-Xmx100g" GenotypeGVCFs -R $Reference -V gendb://GenDB -O oreo_genotype.g.vcf.gz

Filtering:

```
bcftools filter -G 5 -e 'TYPE != "snp" || ALT="*"' -Oz -o oreo_genotype_snp_G5.g.vcf.gz oreo_genotype.g.vcf.gz
bcftools index oreo_genotype_snp_G5.g.vcf.gz
tabix oreo_genotype_snp_G5.g.vcf.gz
gatk VariantFiltration \
  -R $ReferenceO \
  -V oreo_genotype_snp_G5.g.vcf.gz \
  -O oreo_nucrawfiltersnps.vcf.gz \
  --filter-expression "QD < 2.0 || FS > 10.0 || MQ < 30.0 || MQRankSum < -2.0 || ReadPosRankSum < -2.0 || SOR > 3.0 || DP > 180.0" \
  --filter-name "filter" \
  --genotype-filter-expression "DP < 3.0" \
  --genotype-filter-name "lowCov" \
  --set-filtered-genotype-to-no-call true
bcftools view -e 'FILTER!="PASS"' -Oz -o oreo_nucfiltersnps.vcf.gz oreo_nucrawfiltersnps.vcf.gz.gz
```

Phylogeny reconstruction using RAXML (using full sequence mtDNA data):

raxmlHPC-PTHREADS-AVX2 -s Mitodenovo.aln -x $RANDOM -p $RANDOM -f a -# 200 -n Mito -m GTRGAMMA -T 4

Phylogeny reconstruction with RAXML using ascertainment bias correction for SNP data (note that heterozygotes were removed using BCFtools tools first):

raxmlHPC-PTHREADS-AVX2 -s raxml.min4.phy -x $RANDOM -p $RANDOM -f a -# 200 -n test -m ASC_GTRGAMMA --asc-corr=lewis -T 16.support nucSNPs.raxml.support_backup

Pruning for linkage disequilibrium in PLINK:

plink2 --vcf oreo_nucfiltersnps_biallelic_lg.vcf.gz --indep-pairwise 50 10 0.2 --out oreo_ldfilter --allow-extra-chr --set-all-var-ids @:#

plink2 --vcf oreo_nucfiltersnps_biallelic_lg.vcf.gz --extract oreo_ldfilter.prune.in --make-bed --out oreo_pruned --allow-extra-chr --set-all-var-ids @:#

Run ADMIXTURE for K=1-6 and PCA

plink2 --pca 20 --out pruned_pca --bed oreo_pruned.bed --bim oreo_pruned.bim --fam oreo_pruned.fam --allow-extra-chr

cut -f1 oreo_pruned.bim | sort -u | while read line; do counter=$((counter + 1)); echo $line $counter; sed -i "s/${line}/${counter}/g" oreo_pruned.bim ; done

for k in {1..6}; do echo $k; admixture --cv oreo_pruned.bed $k > ${k}.o 2> ${k}.e; done

Table S1. Whole genome resequencing sample details and sequencing statistics.

|  |  |  |  |  | **Sequencing statistics: reads aligned to *O. niloticus*** | | | |
| --- | --- | --- | --- | --- | --- | --- | --- | --- |
| **Sample sequencing name** | **Sample name** | **Species** | **Collection Location** | **Paired reads (n)** | **Mapped (%)** | **Paired (%)** | **Mean coverage (X)** | **>5x coverage (% of genome)** |
| 1657_LIB19618_LDI16937_GGCTAC_L005 | T2J5 | *O.urolepis* | Ligongwe Utete (Rufiji) | 24,395,019 | 98.92 | 88.76 | 5.83 | 59.40 |
| 1657_LIB19643_LDI16962_CGATGT_L008 | T6A2 | *O.urolepis* | Lower Wami | 22,518,296 | 98.99 | 88.67 | 5.41 | 54.67 |
| 1689_LIB19659_LDI16978_ATGTCA_L002 | U1A1 | *O.niloticus* | Uganda | 20,528,899 | 99.21 | 93.14 | 5.02 | 52.13 |
| 1689_LIB19660_LDI16979_CCGTCC_L002 | U3A3 | *O.niloticus* | Uganda | 23,467,486 | 99.18 | 93.33 | 5.74 | 60.83 |
| 1720_LIB20174_LDI17702_TGACCA_L001 | 83-2013 | *O.placidus* | Rovuma | 20,921,460 | 98.75 | 88.09 | 4.97 | 47.81 |
| 1720_LIB20175_LDI17703_CAGATC_L001 | 120-2013 | *O.placidus* | Rovuma | 22,595,615 | 98.94 | 90.46 | 5.45 | 55.80 |
| 1720_LIB20179_LDI17707_CACGAT_L001 | T3J2 | *O.korogwe-N* | Nambawala, Lindi | 21,822,242 | 98.89 | 88.65 | 5.21 | 51.51 |
| 1720_LIB20180_LDI17708_CAGGCG_L001 | T3J4 | *O.korogwe-N* | Nambawala, Lindi | 22,517,046 | 98.93 | 88.26 | 5.38 | 53.48 |
| 1720_LIB20181_LDI17709_CATGGC_L001 | T3J6 | *O.korogwe-N* | Nambawala, Lindi | 22,700,204 | 98.99 | 88.88 | 5.43 | 53.90 |
| 1720_LIB20191_LDI17719_CATGGC_L002 | P4A10 | *O.korogwe-M* | Mlingano Dam | 21,088,993 | 98.27 | 87.63 | 5.01 | 48.55 |
| 1720_LIB20192_LDI17720_CGGAAT_L002 | P4B1 | *O.korogwe-M* | Mlingano Dam | 20,979,050 | 97.39 | 86.48 | 4.93 | 47.64 |
| 1720_LIB20193_LDI17721_TCGGCA_L002 | P4B2 | *O.korogwe-M* | Mlingano Dam | 21,566,100 | 98.11 | 87.25 | 5.10 | 49.34 |
|  |  |  | **MEAN:** | 22,091,701 | 98.71 | 89.13 | 5.29 | 52.92 |

Table S2. Samples used in each of the analyses

| **A** | Morphological analysis of Southern populations |
| --- | --- |
| **B** | Microsatellite analysis of Southern populations |
| **C** | Traditional morphological analysis North and South O. korogwe |
| **D** | Geometric analysis North and South O. korogwe |
| **E** | Microsatellite analysis 6 populations |
| **F** | Whole Genome Resequencing |

| **Species** | **Sample Code** | **Collection Site** | **Collection Date** | **Latitude (decimals)** | **Longitude (decimals)** | **Collector(s)** | **A** | **B** | **C** | **D** | **E** | **F** |
| --- | --- | --- | --- | --- | --- | --- | --- | --- | --- | --- | --- | --- |
| *O. korogwe* | korogwe_Mitupa_M177 | Mitupa, Lindi | 21-27_10_2016 | -10.011 | 39.451 | T. Blackwell; B.P. Ngatunga |  | Y |  |  |  |  |
| *O. korogwe* | korogwe_Mitupa_M178 | Mitupa, Lindi | 21-27_10_2016 | -10.011 | 39.451 | T. Blackwell; B.P. Ngatunga |  | Y |  |  |  |  |
| *O. korogwe* | korogwe_Mitupa_M179 | Mitupa, Lindi | 21-27_10_2016 | -10.011 | 39.451 | T. Blackwell; B.P. Ngatunga |  | Y |  |  |  |  |
| *O. korogwe* | korogwe_Mlingano_P4A10 | Mlingano Dam | 18_08_2015 | -5.122 | 38.857 | A. Shechonge; S. Bradbeer; B.P. Ngatunga; M. Genner |  |  | Y | Y | Y | Y |
| *O. korogwe* | korogwe_Mlingano_P4A6 | Mlingano Dam | 18_08_2015 | -5.122 | 38.857 | A. Shechonge; S. Bradbeer; B.P. Ngatunga; M. Genner |  |  | Y | Y | Y |  |
| *O. korogwe* | korogwe_Mlingano_P4A7 | Mlingano Dam | 18_08_2015 | -5.122 | 38.857 | A. Shechonge; S. Bradbeer; B.P. Ngatunga; M. Genner |  |  | Y | Y | Y |  |
| *O. korogwe* | korogwe_Mlingano_P4A8 | Mlingano Dam | 18_08_2015 | -5.122 | 38.857 | A. Shechonge; S. Bradbeer; B.P. Ngatunga; M. Genner |  |  | Y | Y | Y |  |
| *O. korogwe* | korogwe_Mlingano_P4A9 | Mlingano Dam | 18_08_2015 | -5.122 | 38.857 | A. Shechonge; S. Bradbeer; B.P. Ngatunga; M. Genner |  |  | Y | Y | Y |  |
| *O. korogwe* | korogwe_Mlingano_P4B1 | Mlingano Dam | 18_08_2015 | -5.122 | 38.857 | A. Shechonge; S. Bradbeer; B.P. Ngatunga; M. Genner |  |  | Y | Y | Y | Y |
| *O. korogwe* | korogwe_Mlingano_P4B10 | Mlingano Dam | 18_08_2015 | -5.122 | 38.857 | A. Shechonge; S. Bradbeer; B.P. Ngatunga; M. Genner |  |  | Y | Y | Y |  |
| *O. korogwe* | korogwe_Mlingano_P4B2 | Mlingano Dam | 18_08_2015 | -5.122 | 38.857 | A. Shechonge; S. Bradbeer; B.P. Ngatunga; M. Genner |  |  |  | Y | Y | Y |
| *O. korogwe* | korogwe_Mlingano_P4B3 | Mlingano Dam | 18_08_2015 | -5.122 | 38.857 | A. Shechonge; S. Bradbeer; B.P. Ngatunga; M. Genner |  |  | Y | Y | Y |  |
| *O. korogwe* | korogwe_Mlingano_P4B4 | Mlingano Dam | 18_08_2015 | -5.122 | 38.857 | A. Shechonge; S. Bradbeer; B.P. Ngatunga; M. Genner |  |  |  | Y | Y |  |
| *O. korogwe* | korogwe_Mlingano_P4B5 | Mlingano Dam | 18_08_2015 | -5.122 | 38.857 | A. Shechonge; S. Bradbeer; B.P. Ngatunga; M. Genner |  |  | Y | Y | Y |  |
| *O. korogwe* | korogwe_Mlingano_P4B6 | Mlingano Dam | 18_08_2015 | -5.122 | 38.857 | A. Shechonge; S. Bradbeer; B.P. Ngatunga; M. Genner |  |  | Y | Y | Y |  |
| *O. korogwe* | korogwe_Mlingano_P4B7 | Mlingano Dam | 18_08_2015 | -5.122 | 38.857 | A. Shechonge; S. Bradbeer; B.P. Ngatunga; M. Genner |  |  | Y | Y | Y |  |
| *O. korogwe* | korogwe_Mlingano_P4B8 | Mlingano Dam | 18_08_2015 | -5.122 | 38.857 | A. Shechonge; S. Bradbeer; B.P. Ngatunga; M. Genner |  |  | Y | Y | Y |  |
| *O. korogwe* | korogwe_Mlingano_P4B9 | Mlingano Dam | 18_08_2015 | -5.122 | 38.857 | A. Shechonge; S. Bradbeer; B.P. Ngatunga; M. Genner |  |  | Y | Y | Y |  |
| *O. korogwe* | korogwe_Mlingano_P4C1 | Mlingano Dam | 18_08_2015 | -5.122 | 38.857 | A. Shechonge; S. Bradbeer; B.P. Ngatunga; M. Genner |  |  | Y | Y | Y |  |
| *O. korogwe* | korogwe_Mlingano_P4C2 | Mlingano Dam | 18_08_2015 | -5.122 | 38.857 | A. Shechonge; S. Bradbeer; B.P. Ngatunga; M. Genner |  |  | Y | Y | Y |  |
| *O. korogwe* | korogwe_Mlingano_P4C3 | Mlingano Dam | 18_08_2015 | -5.122 | 38.857 | A. Shechonge; S. Bradbeer; B.P. Ngatunga; M. Genner |  |  | Y | Y | Y |  |
| *O. korogwe* | korogwe_Mlingano_P4C4 | Mlingano Dam | 18_08_2015 | -5.122 | 38.857 | A. Shechonge; S. Bradbeer; B.P. Ngatunga; M. Genner |  |  | Y | Y | Y |  |
| *O. korogwe* | korogwe_Mlingano_P4C5 | Mlingano Dam | 18_08_2015 | -5.122 | 38.857 | A. Shechonge; S. Bradbeer; B.P. Ngatunga; M. Genner |  |  | Y | Y | Y |  |
| *O. korogwe* | korogwe_Mlingano_P4C6 | Mlingano Dam | 18_08_2015 | -5.122 | 38.857 | A. Shechonge; S. Bradbeer; B.P. Ngatunga; M. Genner |  |  |  | Y | Y |  |
| *O. korogwe* | korogwe_Mlingano_P5A10 | Mlingano Dam | 18_08_2015 | -5.122 | 38.857 | A. Shechonge; S. Bradbeer; B.P. Ngatunga; M. Genner |  |  | Y | Y | Y |  |
| *O. korogwe* | korogwe_Mlingano_P5B1 | Mlingano Dam | 18_08_2015 | -5.122 | 38.857 | A. Shechonge; S. Bradbeer; B.P. Ngatunga; M. Genner |  |  | Y | Y | Y |  |
| *O. korogwe* | korogwe_Mlingano_P5B10 | Mlingano Dam | 18_08_2015 | -5.122 | 38.857 | A. Shechonge; S. Bradbeer; B.P. Ngatunga; M. Genner |  |  |  | Y | Y |  |
| *O. korogwe* | korogwe_Mlingano_P5B2 | Mlingano Dam | 18_08_2015 | -5.122 | 38.857 | A. Shechonge; S. Bradbeer; B.P. Ngatunga; M. Genner |  |  |  | Y | Y |  |
| *O. korogwe* | korogwe_Mlingano_P5B3 | Mlingano Dam | 18_08_2015 | -5.122 | 38.857 | A. Shechonge; S. Bradbeer; B.P. Ngatunga; M. Genner |  |  | Y | Y | Y |  |
| *O. korogwe* | korogwe_Mlingano_P5B4 | Mlingano Dam | 18_08_2015 | -5.122 | 38.857 | A. Shechonge; S. Bradbeer; B.P. Ngatunga; M. Genner |  |  | Y | Y | Y |  |
| *O. korogwe* | korogwe_Mlingano_P5B5 | Mlingano Dam | 18_08_2015 | -5.122 | 38.857 | A. Shechonge; S. Bradbeer; B.P. Ngatunga; M. Genner |  |  | Y | Y | Y |  |
| *O. korogwe* | korogwe_Mlingano_P5B6 | Mlingano Dam | 18_08_2015 | -5.122 | 38.857 | A. Shechonge; S. Bradbeer; B.P. Ngatunga; M. Genner |  |  | Y | Y | Y |  |
| *O. korogwe* | korogwe_Mlingano_P5B7 | Mlingano Dam | 18_08_2015 | -5.122 | 38.857 | A. Shechonge; S. Bradbeer; B.P. Ngatunga; M. Genner |  |  | Y | Y | Y |  |
| *O. korogwe* | korogwe_Mlingano_P5B8 | Mlingano Dam | 18_08_2015 | -5.122 | 38.857 | A. Shechonge; S. Bradbeer; B.P. Ngatunga; M. Genner |  |  | Y | Y | Y |  |
| *O. korogwe* | korogwe_Mlingano_P5B9 | Mlingano Dam | 18_08_2015 | -5.122 | 38.857 | A. Shechonge; S. Bradbeer; B.P. Ngatunga; M. Genner |  |  | Y | Y | Y |  |
| *O. korogwe* | korogwe_Mlingano_P5C1 | Mlingano Dam | 18_08_2015 | -5.122 | 38.857 | A. Shechonge; S. Bradbeer; B.P. Ngatunga; M. Genner |  |  | Y | Y | Y |  |
| *O. korogwe* | korogwe_Mlingano_P5C2 | Mlingano Dam | 18_08_2015 | -5.122 | 38.857 | A. Shechonge; S. Bradbeer; B.P. Ngatunga; M. Genner |  |  |  | Y | Y |  |
| *O. korogwe* | korogwe_Mlingano_P5C3 | Mlingano Dam | 18_08_2015 | -5.122 | 38.857 | A. Shechonge; S. Bradbeer; B.P. Ngatunga; M. Genner |  |  | Y | Y | Y |  |
| *O. korogwe* | korogwe_Mlingano_P5C4 | Mlingano Dam | 18_08_2015 | -5.122 | 38.857 | A. Shechonge; S. Bradbeer; B.P. Ngatunga; M. Genner |  |  | Y | Y | Y |  |
| *O. korogwe* | korogwe_Mlingano_P5C5 | Mlingano Dam | 18_08_2015 | -5.122 | 38.857 | A. Shechonge; S. Bradbeer; B.P. Ngatunga; M. Genner |  |  | Y | Y | Y |  |
| *O. korogwe* | korogwe_Mlingano_P5C6 | Mlingano Dam | 18_08_2015 | -5.122 | 38.857 | A. Shechonge; S. Bradbeer; B.P. Ngatunga; M. Genner |  |  | Y | Y | Y |  |
| *O. korogwe* | korogwe_Mlingano_P5C7 | Mlingano Dam | 18_08_2015 | -5.122 | 38.857 | A. Shechonge; S. Bradbeer; B.P. Ngatunga; M. Genner |  |  | Y | Y | Y |  |
| *O. korogwe* | korogwe_Mlingano_P5C8 | Mlingano Dam | 18_08_2015 | -5.122 | 38.857 | A. Shechonge; S. Bradbeer; B.P. Ngatunga; M. Genner |  |  | Y | Y | Y |  |
| *O. korogwe* | korogwe_Nambawala_N63 | Nambawala, Lindi | 21-27_10_2016 | -10.044 | 39.453 | T. Blackwell; B.P. Ngatunga | Y | Y | Y | Y | Y |  |
| *O. korogwe* | korogwe_Nambawala_N66 | Nambawala, Lindi | 21-27_10_2016 | -10.044 | 39.453 | T. Blackwell; B.P. Ngatunga | Y | Y | Y | Y | Y |  |
| *O. korogwe* | korogwe_Nambawala_N71 | Nambawala, Lindi | 21-27_10_2016 | -10.044 | 39.453 | T. Blackwell; B.P. Ngatunga | Y | Y | Y | Y | Y |  |
| *O. korogwe* | korogwe_Nambawala_N75 | Nambawala, Lindi | 21-27_10_2016 | -10.044 | 39.453 | T. Blackwell; B.P. Ngatunga | Y | Y | Y | Y | Y |  |
| *O. korogwe* | korogwe_Nambawala_T3J2 | Nambawala, Lindi | 03_06_2015 | -10.044 | 39.453 | A.Shechonge |  |  |  |  |  | Y |
| *O. korogwe* | korogwe_Nambawala_T3J4 | Nambawala, Lindi | 03_06_2015 | -10.044 | 39.453 | A.Shechonge |  |  |  |  |  | Y |
| *O. korogwe* | korogwe_Nambawala_T3J6 | Nambawala, Lindi | 03_06_2015 | -10.044 | 39.453 | A.Shechonge |  |  |  |  |  | Y |
| *O. korogwe* | korogwe_Nambawala_T4A6 (=TXA6) | Nambawala, Lindi | 03_06_2015 | -10.044 | 39.453 | A.Shechonge |  | Y |  | Y |  |  |
| *O. korogwe* | korogwe_Nambawala_T4A7 (=TXA7) | Nambawala, Lindi | 03_06_2015 | -10.044 | 39.453 | A.Shechonge | Y | Y | Y | Y |  |  |
| *O. korogwe* | korogwe_Nambawala_T4A9 (=TXA9) | Nambawala, Lindi | 03_06_2015 | -10.044 | 39.453 | A.Shechonge | Y | Y | Y | Y |  |  |
| *O. korogwe* | korogwe_Nambawala_T4C1 (=TXC1) | Nambawala, Lindi | 03_06_2015 | -10.044 | 39.453 | A.Shechonge | Y | Y | Y | Y |  |  |
| *O. korogwe* | korogwe_Nambawala_T4C2 (=TXC2) | Nambawala, Lindi | 03_06_2015 | -10.044 | 39.453 | A.Shechonge | Y |  | Y | Y |  |  |
| *O. korogwe* | korogwe_Nambawala_T4C4 (=TXC4) | Nambawala, Lindi | 03_06_2015 | -10.044 | 39.453 | A.Shechonge | Y | Y | Y | Y |  |  |
| *O. korogwe* | korogwe_Rutamba_33A | Rutamba, Lindi | 14_08_2013 | -10.032 | 39.46 | A. Shechonge; B.P. Ngatunga; M. Genner |  | Y |  |  |  |  |
| *O. korogwe* | korogwe_Rutamba_33B | Rutamba, Lindi | 14_08_2013 | -10.032 | 39.46 | A. Shechonge; B.P. Ngatunga; M. Genner |  | Y |  |  |  |  |
| *O. korogwe* | korogwe_Rutamba_33C | Rutamba, Lindi | 14_08_2013 | -10.032 | 39.46 | A. Shechonge; B.P. Ngatunga; M. Genner |  | Y |  |  |  |  |
| *O. korogwe* | korogwe_Rutamba_34B | Rutamba, Lindi | 14_08_2013 | -10.032 | 39.46 | A. Shechonge; B.P. Ngatunga; M. Genner |  | Y |  |  |  |  |
| *O. korogwe* | korogwe_Rutamba_34F | Rutamba, Lindi | 14_08_2013 | -10.032 | 39.46 | A. Shechonge; B.P. Ngatunga; M. Genner |  | Y |  |  |  |  |
| *O. korogwe* | korogwe_Rutamba_42C | Rutamba, Lindi | 14_08_2013 | -10.032 | 39.46 | A. Shechonge; B.P. Ngatunga; M. Genner |  | Y |  |  | Y |  |
| *O. korogwe* | korogwe_Rutamba_43B | Rutamba, Lindi | 14_08_2013 | -10.032 | 39.46 | A. Shechonge; B.P. Ngatunga; M. Genner |  | Y |  |  | Y |  |
| *O. korogwe* | korogwe_Rutamba_43C | Rutamba, Lindi | 14_08_2013 | -10.032 | 39.46 | A. Shechonge; B.P. Ngatunga; M. Genner |  | Y |  |  | Y |  |
| *O. korogwe* | korogwe_Rutamba_R17 | Rutamba, Lindi | 21-27_10_2016 | -10.032 | 39.46 | T. Blackwell; B.P. Ngatunga | Y | Y | Y | Y | Y |  |
| *O. korogwe* | korogwe_Rutamba_R207 | Rutamba, Lindi | 21-27_10_2016 | -10.032 | 39.46 | T. Blackwell; B.P. Ngatunga | Y | Y | Y | Y | Y |  |
| *O. korogwe* | korogwe_Rutamba_R210 | Rutamba, Lindi | 21-27_10_2016 | -10.032 | 39.46 | T. Blackwell; B.P. Ngatunga | Y | Y | Y | Y | Y |  |
| *O. korogwe* | korogwe_Rutamba_R215 | Rutamba, Lindi | 21-27_10_2016 | -10.032 | 39.46 | T. Blackwell; B.P. Ngatunga | Y | Y | Y | Y | Y |  |
| *O. korogwe* | korogwe_Rutamba_R22 | Rutamba, Lindi | 21-27_10_2016 | -10.032 | 39.46 | T. Blackwell; B.P. Ngatunga |  | Y |  |  |  |  |
| *O. korogwe* | korogwe_Rutamba_R222 | Rutamba, Lindi | 21-27_10_2016 | -10.032 | 39.46 | T. Blackwell; B.P. Ngatunga | Y | Y | Y | Y | Y |  |
| *O. korogwe* | korogwe_Rutamba_R23 | Rutamba, Lindi | 21-27_10_2016 | -10.032 | 39.46 | T. Blackwell; B.P. Ngatunga | Y | Y | Y | Y | Y |  |
| *O. korogwe* | korogwe_Rutamba_R240 | Rutamba, Lindi | 21-27_10_2016 | -10.032 | 39.46 | T. Blackwell; B.P. Ngatunga | Y | Y | Y | Y | Y |  |
| *O. korogwe* | korogwe_Rutamba_R240 | Rutamba, Lindi | 21-27_10_2016 | -10.032 | 39.46 | T. Blackwell; B.P. Ngatunga | Y | Y | Y | Y | Y |  |
| *O. korogwe* | korogwe_Rutamba_R244 | Rutamba, Lindi | 21-27_10_2016 | -10.032 | 39.46 | T. Blackwell; B.P. Ngatunga |  | Y | Y |  | Y |  |
| *O. korogwe* | korogwe_Rutamba_R244 | Rutamba, Lindi | 21-27_10_2016 | -10.032 | 39.46 | T. Blackwell; B.P. Ngatunga |  | Y | Y |  | Y |  |
| *O. korogwe* | korogwe_Rutamba_R246 | Rutamba, Lindi | 21-27_10_2016 | -10.032 | 39.46 | T. Blackwell; B.P. Ngatunga | Y | Y | Y | Y | Y |  |
| *O. korogwe* | korogwe_Rutamba_R247 | Rutamba, Lindi | 21-27_10_2016 | -10.032 | 39.46 | T. Blackwell; B.P. Ngatunga | Y | Y | Y | Y | Y |  |
| *O. korogwe* | korogwe_Rutamba_R82 | Rutamba, Lindi | 21-27_10_2016 | -10.032 | 39.46 | T. Blackwell; B.P. Ngatunga |  | Y | Y |  | Y |  |
| *O. korogwe* | korogwe_Rutamba_R83 | Rutamba, Lindi | 21-27_10_2016 | -10.032 | 39.46 | T. Blackwell; B.P. Ngatunga |  | Y | Y |  | Y |  |
| *O. korogwe* | korogwe_Rutamba_R86 | Rutamba, Lindi | 21-27_10_2016 | -10.032 | 39.46 | T. Blackwell; B.P. Ngatunga |  | Y | Y |  | Y |  |
| *O. korogwe* | korogwe_Rutamba_R88 | Rutamba, Lindi | 21-27_10_2016 | -10.032 | 39.46 | T. Blackwell; B.P. Ngatunga |  | Y | Y |  | Y |  |
| *O. korogwe* | korogwe_Zigi_P4C10 | Zigi River | 18_08_2015 | -5.042 | 38.898 | A. Shechonge; S. Bradbeer; B.P. Ngatunga; M. Genner |  |  | Y | Y |  |  |
| *O. korogwe* | korogwe_Zigi_P4C7 | Zigi River | 18_08_2015 | -5.042 | 38.898 | A. Shechonge; S. Bradbeer; B.P. Ngatunga; M. Genner |  |  |  | Y |  |  |
| *O. korogwe* | korogwe_Zigi_P4C8 | Zigi River | 18_08_2015 | -5.042 | 38.898 | A. Shechonge; S. Bradbeer; B.P. Ngatunga; M. Genner |  |  | Y | Y |  |  |
| *O. korogwe* | korogwe_Zigi_P4C9 | Zigi River | 18_08_2015 | -5.042 | 38.898 | A. Shechonge; S. Bradbeer; B.P. Ngatunga; M. Genner |  |  |  | Y |  |  |
| *O. korogwe* | korogwe_Zigi_P4D1 | Zigi River | 18_08_2015 | -5.042 | 38.898 | A. Shechonge; S. Bradbeer; B.P. Ngatunga; M. Genner |  |  | Y | Y |  |  |
| *O. korogwe* | korogwe_Zigi_P4D8 | Zigi River | 18_08_2015 | -5.042 | 38.898 | A. Shechonge; S. Bradbeer; B.P. Ngatunga; M. Genner |  |  | Y |  |  |  |
| *O. korogwe* | korogwe_Zigi_S20K01 | Zigi River | 18_08_2015 | -5.042 | 38.898 | A. Shechonge; S. Bradbeer; B.P. Ngatunga; M. Genner |  |  | Y | Y |  |  |
| *O. korogwe* | korogwe_Zigi_S20K010 | Zigi River | 18_08_2015 | -5.042 | 38.898 | A. Shechonge; S. Bradbeer; B.P. Ngatunga; M. Genner |  |  | Y | Y |  |  |
| *O. korogwe* | korogwe_Zigi_S20K011 | Zigi River | 18_08_2015 | -5.042 | 38.898 | A. Shechonge; S. Bradbeer; B.P. Ngatunga; M. Genner |  |  | Y | Y |  |  |
| *O. korogwe* | korogwe_Zigi_S20K012 | Zigi River | 18_08_2015 | -5.042 | 38.898 | A. Shechonge; S. Bradbeer; B.P. Ngatunga; M. Genner |  |  | Y | Y |  |  |
| *O. korogwe* | korogwe_Zigi_S20K013 | Zigi River | 18_08_2015 | -5.042 | 38.898 | A. Shechonge; S. Bradbeer; B.P. Ngatunga; M. Genner |  |  | Y | Y |  |  |
| *O. korogwe* | korogwe_Zigi_S20K014 | Zigi River | 18_08_2015 | -5.042 | 38.898 | A. Shechonge; S. Bradbeer; B.P. Ngatunga; M. Genner |  |  |  | Y |  |  |
| *O. korogwe* | korogwe_Zigi_S20K015 | Zigi River | 18_08_2015 | -5.042 | 38.898 | A. Shechonge; S. Bradbeer; B.P. Ngatunga; M. Genner |  |  |  | Y |  |  |
| *O. korogwe* | korogwe_Zigi_S20K016 | Zigi River | 18_08_2015 | -5.042 | 38.898 | A. Shechonge; S. Bradbeer; B.P. Ngatunga; M. Genner |  |  |  | Y |  |  |
| *O. korogwe* | korogwe_Zigi_S20K017 | Zigi River | 18_08_2015 | -5.042 | 38.898 | A. Shechonge; S. Bradbeer; B.P. Ngatunga; M. Genner |  |  | Y | Y |  |  |
| *O. korogwe* | korogwe_Zigi_S20K018 | Zigi River | 18_08_2015 | -5.042 | 38.898 | A. Shechonge; S. Bradbeer; B.P. Ngatunga; M. Genner |  |  | Y | Y |  |  |
| *O. korogwe* | korogwe_Zigi_S20K019 | Zigi River | 18_08_2015 | -5.042 | 38.898 | A. Shechonge; S. Bradbeer; B.P. Ngatunga; M. Genner |  |  | Y | Y |  |  |
| *O. korogwe* | korogwe_Zigi_S20K02 | Zigi River | 18_08_2015 | -5.042 | 38.898 | A. Shechonge; S. Bradbeer; B.P. Ngatunga; M. Genner |  |  | Y | Y |  |  |
| *O. korogwe* | korogwe_Zigi_S20K020 | Zigi River | 18_08_2015 | -5.042 | 38.898 | A. Shechonge; S. Bradbeer; B.P. Ngatunga; M. Genner |  |  | Y | Y |  |  |
| *O. korogwe* | korogwe_Zigi_S20K021 | Zigi River | 18_08_2015 | -5.042 | 38.898 | A. Shechonge; S. Bradbeer; B.P. Ngatunga; M. Genner |  |  | Y | Y |  |  |
| *O. korogwe* | korogwe_Zigi_S20K022 | Zigi River | 18_08_2015 | -5.042 | 38.898 | A. Shechonge; S. Bradbeer; B.P. Ngatunga; M. Genner |  |  |  | Y |  |  |
| *O. korogwe* | korogwe_Zigi_S20K023 | Zigi River | 18_08_2015 | -5.042 | 38.898 | A. Shechonge; S. Bradbeer; B.P. Ngatunga; M. Genner |  |  | Y | Y |  |  |
| *O. korogwe* | korogwe_Zigi_S20K024 | Zigi River | 18_08_2015 | -5.042 | 38.898 | A. Shechonge; S. Bradbeer; B.P. Ngatunga; M. Genner |  |  | Y | Y |  |  |
| *O. korogwe* | korogwe_Zigi_S20K03 | Zigi River | 18_08_2015 | -5.042 | 38.898 | A. Shechonge; S. Bradbeer; B.P. Ngatunga; M. Genner |  |  | Y | Y |  |  |
| *O. korogwe* | korogwe_Zigi_S20K04 | Zigi River | 18_08_2015 | -5.042 | 38.898 | A. Shechonge; S. Bradbeer; B.P. Ngatunga; M. Genner |  |  | Y | Y |  |  |
| *O. korogwe* | korogwe_Zigi_S20K05 | Zigi River | 18_08_2015 | -5.042 | 38.898 | A. Shechonge; S. Bradbeer; B.P. Ngatunga; M. Genner |  |  |  | Y |  |  |
| *O. korogwe* | korogwe_Zigi_S20K06 | Zigi River | 18_08_2015 | -5.042 | 38.898 | A. Shechonge; S. Bradbeer; B.P. Ngatunga; M. Genner |  |  | Y | Y |  |  |
| *O. korogwe* | korogwe_Zigi_S20K07 | Zigi River | 18_08_2015 | -5.042 | 38.898 | A. Shechonge; S. Bradbeer; B.P. Ngatunga; M. Genner |  |  | Y | Y |  |  |
| *O. korogwe* | korogwe_Zigi_S20K08 | Zigi River | 18_08_2015 | -5.042 | 38.898 | A. Shechonge; S. Bradbeer; B.P. Ngatunga; M. Genner |  |  | Y | Y |  |  |
| *O. korogwe* | korogwe_Zigi_S20K09 | Zigi River | 18_08_2015 | -5.042 | 38.898 | A. Shechonge; S. Bradbeer; B.P. Ngatunga; M. Genner |  |  |  | Y |  |  |
| *O. korogwe* | korogwe_Zigi_Z10 | Zigi River | 18_08_2015 | -5.042 | 38.898 | A. Shechonge; S. Bradbeer; B.P. Ngatunga; M. Genner |  |  |  |  | Y |  |
| *O. korogwe* | korogwe_Zigi_Z12 | Zigi River | 18_08_2015 | -5.042 | 38.898 | A. Shechonge; S. Bradbeer; B.P. Ngatunga; M. Genner |  |  |  |  | Y |  |
| *O. korogwe* | korogwe_Zigi_Z17 | Zigi River | 18_08_2015 | -5.042 | 38.898 | A. Shechonge; S. Bradbeer; B.P. Ngatunga; M. Genner |  |  |  |  | Y |  |
| *O. korogwe* | korogwe_Zigi_Z19 | Zigi River | 18_08_2015 | -5.042 | 38.898 | A. Shechonge; S. Bradbeer; B.P. Ngatunga; M. Genner |  |  |  |  | Y |  |
| *O. korogwe* | korogwe_Zigi_Z2 | Zigi River | 18_08_2015 | -5.042 | 38.898 | A. Shechonge; S. Bradbeer; B.P. Ngatunga; M. Genner |  |  |  |  | Y |  |
| *O. korogwe* | korogwe_Zigi_Z20 | Zigi River | 18_08_2015 | -5.042 | 38.898 | A. Shechonge; S. Bradbeer; B.P. Ngatunga; M. Genner |  |  |  |  | Y |  |
| *O. korogwe* | korogwe_Zigi_Z21 | Zigi River | 18_08_2015 | -5.042 | 38.898 | A. Shechonge; S. Bradbeer; B.P. Ngatunga; M. Genner |  |  |  |  | Y |  |
| *O. korogwe* | korogwe_Zigi_Z22 | Zigi River | 18_08_2015 | -5.042 | 38.898 | A. Shechonge; S. Bradbeer; B.P. Ngatunga; M. Genner |  |  |  |  | Y |  |
| *O. korogwe* | korogwe_Zigi_Z24 | Zigi River | 18_08_2015 | -5.042 | 38.898 | A. Shechonge; S. Bradbeer; B.P. Ngatunga; M. Genner |  |  |  |  | Y |  |
| *O. korogwe* | korogwe_Zigi_Z3 | Zigi River | 18_08_2015 | -5.042 | 38.898 | A. Shechonge; S. Bradbeer; B.P. Ngatunga; M. Genner |  |  |  |  | Y |  |
| *O. korogwe* | korogwe_Zigi_Z4 | Zigi River | 18_08_2015 | -5.042 | 38.898 | A. Shechonge; S. Bradbeer; B.P. Ngatunga; M. Genner |  |  |  |  | Y |  |
| *O. korogwe* | korogwe_Zigi_Z5 | Zigi River | 18_08_2015 | -5.042 | 38.898 | A. Shechonge; S. Bradbeer; B.P. Ngatunga; M. Genner |  |  |  |  | Y |  |
| *O. korogwe* | korogwe_Zigi_Z6 | Zigi River | 18_08_2015 | -5.042 | 38.898 | A. Shechonge; S. Bradbeer; B.P. Ngatunga; M. Genner |  |  |  |  | Y |  |
| *O. korogwe* | korogwe_Zigi_Z7 | Zigi River | 18_08_2015 | -5.042 | 38.898 | A. Shechonge; S. Bradbeer; B.P. Ngatunga; M. Genner |  |  |  |  | Y |  |
| *O. korogwe* | korogwe_Zigi_Z8 | Zigi River | 18_08_2015 | -5.042 | 38.898 | A. Shechonge; S. Bradbeer; B.P. Ngatunga; M. Genner |  |  |  |  | Y |  |
| *O. korogwe* | korogwe_Zigi_Z9 | Zigi River | 18_08_2015 | -5.042 | 38.898 | A. Shechonge; S. Bradbeer; B.P. Ngatunga; M. Genner |  |  |  |  | Y |  |
| *O. niloticus* | niloticus_Mitupa_M183 | Mitupa, Lindi | 21-27_10_2016 | -10.011 | 39.451 | T. Blackwell; B.P. Ngatunga | Y | Y |  |  |  |  |
| *O. niloticus* | niloticus_Mitupa_M184 | Mitupa, Lindi | 21-27_10_2016 | -10.011 | 39.451 | T. Blackwell; B.P. Ngatunga | Y | Y |  |  |  |  |
| *O. niloticus* | niloticus_Mitupa_M185 | Mitupa, Lindi | 21-27_10_2016 | -10.011 | 39.451 | T. Blackwell; B.P. Ngatunga | Y | Y |  |  |  |  |
| *O. niloticus* | niloticus_Nambawala_N165 | Nambawala, Lindi | 21-27_10_2016 | -10.044 | 39.453 | T. Blackwell; B.P. Ngatunga |  | Y |  |  |  |  |
| *O. niloticus* | niloticus_Nambawala_N166 | Nambawala, Lindi | 21-27_10_2016 | -10.044 | 39.453 | T. Blackwell; B.P. Ngatunga |  | Y |  |  |  |  |
| *O. niloticus* | niloticus_Nambawala_N167 | Nambawala, Lindi | 21-27_10_2016 | -10.044 | 39.453 | T. Blackwell; B.P. Ngatunga |  | Y |  |  |  |  |
| *O. niloticus* | niloticus_Nambawala_N170 | Nambawala, Lindi | 21-27_10_2016 | -10.044 | 39.453 | T. Blackwell; B.P. Ngatunga |  | Y |  |  |  |  |
| *O. niloticus* | niloticus_Nambawala_N171 | Nambawala, Lindi | 21-27_10_2016 | -10.044 | 39.453 | T. Blackwell; B.P. Ngatunga |  | Y |  |  |  |  |
| *O. niloticus* | niloticus_Nambawala_N172 | Nambawala, Lindi | 21-27_10_2016 | -10.044 | 39.453 | T. Blackwell; B.P. Ngatunga | Y | Y |  |  |  |  |
| *O. niloticus* | niloticus_Nambawala_T4C3 (=TXC3) | Nambawala, Lindi | 03_06_2015 | -10.044 | 39.453 | A.Shechonge |  | Y |  |  |  |  |
| *O. niloticus* | niloticus_Rutamba_R13 | Rutamba, Lindi | 21-27_10_2016 | -10.032 | 39.46 | T. Blackwell; B.P. Ngatunga | Y | Y |  |  |  |  |
| *O. niloticus* | niloticus_Rutamba_R14 | Rutamba, Lindi | 21-27_10_2016 | -10.032 | 39.46 | T. Blackwell; B.P. Ngatunga | Y | Y |  |  |  |  |
| *O. niloticus* | niloticus_Rutamba_R152 | Rutamba, Lindi | 21-27_10_2016 | -10.032 | 39.46 | T. Blackwell; B.P. Ngatunga | Y | Y |  |  |  |  |
| *O. niloticus* | niloticus_Rutamba_R165 | Rutamba, Lindi | 21-27_10_2016 | -10.032 | 39.46 | T. Blackwell; B.P. Ngatunga | Y |  |  |  |  |  |
| *O. niloticus* | niloticus_Rutamba_R167 | Rutamba, Lindi | 21-27_10_2016 | -10.032 | 39.46 | T. Blackwell; B.P. Ngatunga | Y |  |  |  |  |  |
| *O. niloticus* | niloticus_Rutamba_R171 | Rutamba, Lindi | 21-27_10_2016 | -10.032 | 39.46 | T. Blackwell; B.P. Ngatunga | Y |  |  |  |  |  |
| *O. niloticus* | niloticus_Rutamba_R18 | Rutamba, Lindi | 21-27_10_2016 | -10.032 | 39.46 | T. Blackwell; B.P. Ngatunga |  | Y |  |  |  |  |
| *O. niloticus* | niloticus_Rutamba_R223 | Rutamba, Lindi | 21-27_10_2016 | -10.032 | 39.46 | T. Blackwell; B.P. Ngatunga | Y | Y |  |  |  |  |
| *O. niloticus* | niloticus_Rutamba_R225 | Rutamba, Lindi | 21-27_10_2016 | -10.032 | 39.46 | T. Blackwell; B.P. Ngatunga | Y | Y |  |  |  |  |
| *O. niloticus* | niloticus_Rutamba_R29 | Rutamba, Lindi | 21-27_10_2016 | -10.032 | 39.46 | T. Blackwell; B.P. Ngatunga | Y | Y |  |  |  |  |
| *O. niloticus* | niloticus_Uganda_U1A1 | Uganda | 29_10_2015 | - | - | N. Kazosi |  |  |  |  |  | Y |
| *O. niloticus* | niloticus_Uganda_U3A3 | Uganda | 29_10_2015 | - | - | N. Kazosi |  |  |  |  |  | Y |
| *O. placidus* | placidus_Chidya_142A | Lake Chidya | 18_08_2013 | -10.597 | 40.155 | A. Shechonge; B.P. Ngatunga; M. Genner |  |  |  |  | Y |  |
| *O. placidus* | placidus_Chidya_142B | Lake Chidya | 18_08_2013 | -10.597 | 40.155 | A. Shechonge; B.P. Ngatunga; M. Genner |  |  |  |  | Y |  |
| *O. placidus* | placidus_Chidya_142D | Lake Chidya | 18_08_2013 | -10.597 | 40.155 | A. Shechonge; B.P. Ngatunga; M. Genner |  |  |  |  | Y |  |
| *O. placidus* | placidus_Chidya_143A | Lake Chidya | 18_08_2013 | -10.597 | 40.155 | A. Shechonge; B.P. Ngatunga; M. Genner |  |  |  |  | Y |  |
| *O. placidus* | placidus_Chidya_143C | Lake Chidya | 18_08_2013 | -10.597 | 40.155 | A. Shechonge; B.P. Ngatunga; M. Genner |  |  |  |  | Y |  |
| *O. placidus* | placidus_Chidya_143D | Lake Chidya | 18_08_2013 | -10.597 | 40.155 | A. Shechonge; B.P. Ngatunga; M. Genner |  |  |  |  | Y |  |
| *O. placidus* | placidus_Chidya_144A | Lake Chidya | 18_08_2013 | -10.597 | 40.155 | A. Shechonge; B.P. Ngatunga; M. Genner |  |  |  |  | Y |  |
| *O. placidus* | placidus_Chidya_144C | Lake Chidya | 18_08_2013 | -10.597 | 40.155 | A. Shechonge; B.P. Ngatunga; M. Genner |  |  |  |  | Y |  |
| *O. placidus* | placidus_Chidya_144D | Lake Chidya | 18_08_2013 | -10.597 | 40.155 | A. Shechonge; B.P. Ngatunga; M. Genner |  |  |  |  | Y |  |
| *O. placidus* | placidus_Chidya_145C | Lake Chidya | 18_08_2013 | -10.597 | 40.155 | A. Shechonge; B.P. Ngatunga; M. Genner |  |  |  |  | Y |  |
| *O. placidus* | placidus_Rovuma_120-2013 | Muguwesi, Rovuma | 17_08_2013 | -10.847 | 37.474 | A. Shechonge; B.P. Ngatunga; M. Genner |  |  |  |  |  | Y |
| *O. placidus* | placidus_Rovuma_83-2013 | Rovuma River | 16_08_2013 | -11.414 | 38.492 | A. Shechonge; B.P. Ngatunga; M. Genner |  |  |  |  |  | Y |
| *O. urolepis* | urolepis_Utete_T2J05 | Lugongwe Utete | 11_03_2015 | -8 | 38.759 | A. Shechonge; B.P. Ngatunga |  |  |  |  |  | Y |
| *O. urolepis* | urolepis_Utete_T3A01 | Lugongwe Utete | 11_03_2015 | -8 | 38.759 | A. Shechonge; B.P. Ngatunga |  |  |  |  | Y |  |
| *O. urolepis* | urolepis_Utete_T3A02 | Lugongwe Utete | 11_03_2015 | -8 | 38.759 | A. Shechonge; B.P. Ngatunga |  |  |  |  | Y |  |
| *O. urolepis* | urolepis_Utete_T3A03 | Lugongwe Utete | 11_03_2015 | -8 | 38.759 | A. Shechonge; B.P. Ngatunga |  |  |  |  | Y |  |
| *O. urolepis* | urolepis_Utete_T3A04 | Lugongwe Utete | 11_03_2015 | -8 | 38.759 | A. Shechonge; B.P. Ngatunga |  |  |  |  | Y |  |
| *O. urolepis* | urolepis_Utete_T3A05 | Lugongwe Utete | 11_03_2015 | -8 | 38.759 | A. Shechonge; B.P. Ngatunga |  |  |  |  | Y |  |
| *O. urolepis* | urolepis_Utete_T3A06 | Lugongwe Utete | 11_03_2015 | -8 | 38.759 | A. Shechonge; B.P. Ngatunga |  |  |  |  | Y |  |
| *O. urolepis* | urolepis_Utete_T3A07 | Lugongwe Utete | 11_03_2015 | -8 | 38.759 | A. Shechonge; B.P. Ngatunga |  |  |  |  | Y |  |
| *O. urolepis* | urolepis_Utete_T3A08 | Lugongwe Utete | 11_03_2015 | -8 | 38.759 | A. Shechonge; B.P. Ngatunga |  |  |  |  | Y |  |
| *O. urolepis* | urolepis_Utete_T3A09 | Lugongwe Utete | 11_03_2015 | -8 | 38.759 | A. Shechonge; B.P. Ngatunga |  |  |  |  | Y |  |
| *O. urolepis* | urolepis_Utete_T3A10 | Lugongwe Utete | 11_03_2015 | -8 | 38.759 | A. Shechonge; B.P. Ngatunga |  |  |  |  | Y |  |
| *O. urolepis* | urolepis_Utete_T3B01 | Lugongwe Utete | 11_03_2015 | -8 | 38.759 | A. Shechonge; B.P. Ngatunga |  |  |  |  | Y |  |
| *O. urolepis* | urolepis_Utete_T3B02 | Lugongwe Utete | 11_03_2015 | -8 | 38.759 | A. Shechonge; B.P. Ngatunga |  |  |  |  | Y |  |
| *O. urolepis* | urolepis_Utete_T3B03 | Lugongwe Utete | 11_03_2015 | -8 | 38.759 | A. Shechonge; B.P. Ngatunga |  |  |  |  | Y |  |
| *O. urolepis* | urolepis_Utete_T3B04 | Lugongwe Utete | 11_03_2015 | -8 | 38.759 | A. Shechonge; B.P. Ngatunga |  |  |  |  | Y |  |
| *O. urolepis* | urolepis_Utete_T3B05 | Lugongwe Utete | 11_03_2015 | -8 | 38.759 | A. Shechonge; B.P. Ngatunga |  |  |  |  | Y |  |
| *O. urolepis* | urolepis_Utete_T3B06 | Lugongwe Utete | 11_03_2015 | -8 | 38.759 | A. Shechonge; B.P. Ngatunga |  |  |  |  | Y |  |
| *O. urolepis* | urolepis_Utete_T3B07 | Lugongwe Utete | 11_03_2015 | -8 | 38.759 | A. Shechonge; B.P. Ngatunga |  |  |  |  | Y |  |
| *O. urolepis* | urolepis_Utete_T3B08 | Lugongwe Utete | 11_03_2015 | -8 | 38.759 | A. Shechonge; B.P. Ngatunga |  |  |  |  | Y |  |
| *O. urolepis* | urolepis_Utete_T3B09 | Lugongwe Utete | 11_03_2015 | -8 | 38.759 | A. Shechonge; B.P. Ngatunga |  |  |  |  | Y |  |
| *O. urolepis* | urolepis_Utete_T3B10 | Lugongwe Utete | 11_03_2015 | -8 | 38.759 | A. Shechonge; B.P. Ngatunga |  |  |  |  | Y |  |
| *O. urolepis* | urolepis_Utete_T3C01 | Lugongwe Utete | 11_03_2015 | -8 | 38.759 | A. Shechonge; B.P. Ngatunga |  |  |  |  | Y |  |
| *O. urolepis* | urolepis_Utete_T3C02 | Lugongwe Utete | 11_03_2015 | -8 | 38.759 | A. Shechonge; B.P. Ngatunga |  |  |  |  | Y |  |
| *O. urolepis* | urolepis_Utete_T3C03 | Lugongwe Utete | 11_03_2015 | -8 | 38.759 | A. Shechonge; B.P. Ngatunga |  |  |  |  | Y |  |
| *O. urolepis* | urolepis_Utete_T3C04 | Lugongwe Utete | 11_03_2015 | -8 | 38.759 | A. Shechonge; B.P. Ngatunga |  |  |  |  | Y |  |
| *O. urolepis* | urolepis_Utete_T3C05 | Lugongwe Utete | 11_03_2015 | -8 | 38.759 | A. Shechonge; B.P. Ngatunga |  |  |  |  | Y |  |
| *O. urolepis* | urolepis_Utete_T3C06 | Lugongwe Utete | 11_03_2015 | -8 | 38.759 | A. Shechonge; B.P. Ngatunga |  |  |  |  | Y |  |
| *O. urolepis* | urolepis_Wami_T6A02 | Mbuyuni, Wami | 22_07_2015 | -6.25149 | 38.6875 | G. Turner |  |  |  |  |  | Y |
| OKxON hybrid | hybrid_Mitupa_M182 | Mitupa, Lindi | 21-27_10_2016 | -10.011 | 39.451 | T. Blackwell; B.P. Ngatunga | Y | Y |  |  |  |  |
| OKxON hybrid | hybrid_Nambawala_N174 | Nambawala, Lindi | 21-27_10_2016 | -10.044 | 39.453 | T. Blackwell; B.P. Ngatunga | Y | Y |  |  |  |  |
| OKxON hybrid | hybrid_Nambawala_N65 | Nambawala, Lindi | 21-27_10_2016 | -10.044 | 39.453 | T. Blackwell; B.P. Ngatunga | Y | Y |  |  |  |  |
| OKxON hybrid | hybrid_Nambawala_N70 | Nambawala, Lindi | 21-27_10_2016 | -10.044 | 39.453 | T. Blackwell; B.P. Ngatunga | Y | Y |  |  |  |  |
| OKxON hybrid | hybrid_Nambawala_T4A5 (=TXA5) | Nambawala, Lindi | 03_06_2015 | -10.044 | 39.453 | A.Shechonge | Y | Y |  |  |  |  |
| OKxON hybrid | hybrid_Nambawala_T4C5 (=TXC5) | Nambawala, Lindi | 03_06_2015 | -10.044 | 39.453 | A.Shechonge | Y | Y |  |  |  |  |
| OKxON hybrid | hybrid_Rutamba_R144 | Rutamba, Lindi | 21-27_10_2016 | -10.032 | 39.46 | T. Blackwell; B.P. Ngatunga | Y | Y |  |  |  |  |
| OKxON hybrid | hybrid_Rutamba_R245 | Rutamba, Lindi | 21-27_10_2016 | -10.032 | 39.46 | T. Blackwell; B.P. Ngatunga | Y | Y |  |  |  |  |

Table S3: Microsatellite loci primer sequences and sources.

| Marker name | Genbank Accession | Primer sequence (forward) | Primer sequence (reverse) | Motif |
| --- | --- | --- | --- | --- |
| OMO043 | JX204857 | GGGGTCATTCGGTTTATTGGTTAT | AGGGCAGGTCACGGGTTCG | (TTTG)8 |
| OMO093 | JX204891 | AAGCCCCACATAGACGACCAGAGA | CAGAAACGGTGCCTGTTCCAGAA | (CAT)8 |
| OMO100 | JX204895 | CCTTCCCCACCACTACCCTCATAA | CCCGCCCACACCTGACGA | (ATT)18 |
| OMO114 | JX204905 | ACGCCTTAATGCTGCCTTCAAGA | TGATGCTCACCCCGTTCCTCA | (GTT)11 |
| OMO129 | JX204914 | TTGGCAGGCTAAGTACTATTTCAT | GAGCGAATGGTTGTCTGTCTCT | (CCAT)9 |
| OMO161 | JX204924 | ACTTTGACAAAAGAAGTGTAACAA | AGGGGAGGAGAAAATAAACTGTAT | (TAA)10 |
| OMO219 | JX204964 | ATCCCCTTCTTTCCATCCCTGTC | AAGGCCTCTGTGAGCTGATTGATT | (TTTTG)10 |
| OMO229 | JX204973 | GCGACTTTTTCTTTGCACATTTTT | AACTGAACCGCCATCATAATCATC | (GTT)9 |
| OMO248 | JX204987 | AAAGACACAAAGAGAAACTAATCA | GGATGAATATTTAAAATCAGTCAG | (TCA)9 |
| OMO337 | JX205052 | TAGGAGAGGCATAGGTTGTCAAAT | CAAGAGTCTAGGAGGGAATCAAAA | (GTTT)7 |
| OMO361 | JX205069 | TGACAGCGAGCCAGAATGGAAGTA | AAAAGTGAAAGGGGCACAGTGAGG | (CTT)17 |
| OMO391 | GR699257 | AGACATCTGTACGCTCTTTACGAA | AGTGCTAGAGGGAAGGGGCTGTA | (GAT)9 |
| OMO392 | GR698887 | CTGGCTTAACTTCTCTACTGGACA | TCTACTCAAAACTGGCAACAAAAC | (GAATA)7 |
| OMO397 | GR693794 | ACGCGTGTTTGAGATATTTAGATT | GAACAAACAAGGGGAGTGG | (GATT)7 |
| OM-01 | GU391020 | TTTAAAGTTACACAGCAGTACAAAG | TTGTAGCATTTCAACACAGTCTC | (GT)20 |
| OM-03 | GU391022 | CTTTTTAATGAGCAACTTTTAAGTC | TGTGAATTTGACAACTTCCTTTC | (GATA)47 |
| OM-04 | GU391022 | AGCTCAAAACCTCATACAAAGG | GCAGAGATGTCAGATGTTGTTC | (GACA)6 (GATA)16 |
| OM-09 | GU391028 | GGCTACAACACCTGGATGG | TTGGGCTTACTGAAGCTGAC | (GT)26 |

Table S4. Genetic (microsatellite) diversity of the focal populations of *O. korogwe*, *O. urolepis* and *O. placidus*. N - number of individuals, NA -number alleles, Ho - Observed heterozygosity, He - Expected heterozygosity, P - probability of Hardy Weinberg Equilibrium.

| **Site** | **Species** |  | **OMO219** | **OMO229** | **OMO337** | **OMO391** | **OMO392** | **OMO397** | **OMO09** | **OMO043** | **OMO129** | **OMO03** | **OMO04** | **OMO01** | **OMO114** |
| --- | --- | --- | --- | --- | --- | --- | --- | --- | --- | --- | --- | --- | --- | --- | --- |
| Mlingano | *O. korogwe* | N | 33 | - | - | - | 40 | 40 | - | - | - | - | 35 | 35 | 40 |
|  |  | NA | 3 | - | - | - | 3 | 3 | - | - | - | - | 8 | 3 | 2 |
|  |  | Ho | 0.21 | - | - | - | 0.45 | 0.58 | - | - | - | - | 0.83 | 0.71 | 0.33 |
|  |  | He | 0.25 | - | - | - | 0.38 | 0.63 | - | - | - | - | 0.82 | 0.57 | 0.28 |
|  |  | P | 0.03 | - | - | - | 0.6 | 0.45 | - | - | - | - | 0.99 | 0.21 | 0.56 |
| Zigi River | *O. korogwe* | N | 12 | 16 | - | 15 | 9 | 16 | 16 | 16 | - | 5 | - | 14 | 14 |
|  |  | NA | 2 | 3 | - | 4 | 3 | 3 | 2 | 3 | - | 4 | - | 3 | 2 |
|  |  | Ho | 0.17 | 0.19 | - | 0.8 | 0.33 | 0.81 | 0.13 | 0.13 | - | 0 | - | 0.07 | 0 |
|  |  | He | 0.16 | 0.28 | - | 0.65 | 0.54 | 0.66 | 0.31 | 0.23 | - | 0.8 | - | 0.47 | 0.14 |
|  |  | P | 1 | 0.05 | - | 0.82 | 0.17 | 0.5 | 0.05 | 0.19 | - | <0.001 | - | <0.001 | 0.04 |
| Lake Chidya | *O. placidus* | N | 8 | 10 | 10 | 10 | - | 9 | 5 | - | 10 | 10 | 10 | - | - |
|  |  | NA | 2 | 5 | 3 | 2 | - | 4 | 5 | = | 5 | 11 | 5 |  |  |
|  |  | Ho | 0.13 | 0.9 | 0.3 | 0.1 | - | 0.56 | 0.2 | - | 0.7 | 0.3 | 0.7 | - | - |
|  |  | He | 0.13 | 0.73 | 0.43 | 0.1 | - | 0.66 | 0.87 | - | 0.62 | 0.94 | 0.62 | - | - |
|  |  | P | 1 | 0.73 | 0.09 | 1 | - | 0.21 | <0.001 | - | 0.77 | <0.001 | 0.77 | - | - |
| Rufiji | *O. urolepis* | N | 26 | 25 | 26 | 26 | 26 | 26 | 22 | 25 | 26 | 21 | 19 | 22 | 25 |
|  |  | NA | 7 | 8 | 5 | 4 | 3 | 8 | 18 | 4 | 4 | 15 | 18 | 18 | 7 |
|  |  | Ho | 0.77 | 0.84 | 0.27 | 0.54 | 0.35 | 0.77 | 0.86 | 0.52 | 0.31 | 0.52 | 0.74 | 0.73 | 0.84 |
|  |  | He | 0.79 | 0.79 | 0.67 | 0.69 | 0.3 | 0.81 | 0.88 | 0.7 | 0.28 | 0.92 | 0.96 | 0.9 | 0.77 |
|  |  | P | 0.42 | 0.47 | <0.001 | 0.22 | 1 | 0.46 | 0.44 | 0.03 | 1 | <0.001 | <0.01 | 0.05 | 0.6 |
| Rutamba | *O. korogwe* | N | 16 | - | 17 | 17 | 11 | 17 | 16 | - | 17 | 8 | 17 | 13 | 13 |
|  |  | NA | 2 | - | 2 | 2 | 3 | 3 | 2 | - | 2 | 5 | 2 | 5 | 2 |
|  |  | Ho | 0.19 | - | 0 | 0.12 | 0.45 | 0.29 | 0.25 | - | 0 | 0.13 | 0.06 | 0.15 | 0.62 |
|  |  | He | 0.5 | - | 0.11 | 0.11 | 0.65 | 0.27 | 0.31 | - | 0.51 | 0.81 | 0.06 | 0.63 | 0.52 |
|  |  | P | 0.03 | - | 0.03 | 1 | 0.04 | 1 | 0.43 | - | <0.001 | <0.001 | 1 | <0.001 | 0.6 |
| Rutamba | *O. niloticus* | N | 12 | 13 | 13 | - | 12 | 12 | 12 | - | - | 11 | 13 | 6 | 8 |
|  |  | NA | 4 | 2 | 2 | - | 4 | 2 | 3 | - | - | 5 | 2 | 3 | 3 |
|  |  | Ho | 0.08 | 0.54 | 0.08 | - | 0.5 | 0.33 | 0.17 | - | - | 0.82 | 0.08 | 0.17 | 0.38 |
|  |  | He | 0.72 | 0.51 | 0.08 | - | 0.64 | 0.39 | 0.65 | - | - | 0.77 | 0.08 | 0.32 | 0.64 |
|  |  | P | <0.001 | 1 | 1 | - | 0.08 | 1 | <0.001 | - | - | 0.36 | 1 | 0.09 | 0.14 |
| Nambawala | *O. korogwe* | N | 7 | 10 | 10 | - | 4 | 10 | 10 | - | 9 | 1 | 10 | 4 | 4 |
|  |  | NA | 2 | 2 | 2 | - | 3 | 2 | 2 | - | 2 | 2 | 2 | 3 | 2 |
|  |  | Ho | 0.29 | 0.3 | 0 | - | 0.5 | 0.2 | 0.2 | - | 0 | 1 | 0.2 | 0.25 | 0.25 |
|  |  | He | 0.26 | 0.27 | 0.19 | - | 0.61 | 0.19 | 0.34 | - | 0.47 | 1 | 0.19 | 0.61 | 0.25 |
|  |  | P | 1 | 1 | 0.05 | - | 0.43 | 1 | 0.31 | - | <0.001 | 1 | 1 | 0.14 | 1 |
| Nambawala | *O. niloticus* | N | 6 | 6 | - | - | 6 | 6 | 6 | 6 | 6 | 6 | 6 | 6 | 6 |
|  |  | NA | 2 | 2 | - | - | 3 | 3 | 2 | 2 | 2 | 2 | 2 | 6 | 4 |
|  |  | Ho | 0.33 | 0.5 | - | - | 1 | 0.17 | 0.33 | 0.17 | 0.17 | 0.17 | 0.17 | 0.5 | 0.5 |
|  |  | He | 0.48 | 0.53 | - | - | 0.71 | 0.62 | 0.3 | 0.17 | 0.17 | 0.17 | 0.17 | 0.8 | 0.79 |
|  |  | P | 1 | 1 | - | - | 0.58 | 0.03 | 1 | 1 | 1 | 1 | 1 | 0.06 | 0.32 |

Table S5. WGR quality filtering thresholds applied to the SNP and indel datasets.

|  | Nuclear SNP |
| --- | --- |
| QD | < 2.0 |
| FS | > 10.0 |
| SOR | > 3.0 |
| MQ | < 30.0 |
| MQRankSum | < -2.0 |
| ReadPosRankSum | < -2.0 |
| InbreedingCoeff | - |
| DP | > 180.0 |
| Per-Sample DP | < 3.0 |

Filtering parameters (definitions per the GATK website: <https://gatkforums.broadinstitute.org/>)

QD: QualbyDepth - variant confidence divided by the unfiltered depth of non-reference samples.

FS: FisherStrand - Phred-scaled p-value using Fisher’s Exact Test to detect strand bias

SOR: StrandOddsRatio - aims to evaluate whether there is strand bias in the data

MQ: RMSMappingQuality - Root Mean Square of the mapping quality of the reads across all samples

MQRankSum: MappingQualityRankSumTest - The u-based z-approximation from the Mann-Whitney Rank Sum Test for mapping qualities

ReadPosRankSum: ReadPosRankSumTest - the u-based z-approximation from the Mann-Whitney Rank Sum Test for the distance from the end of the read for reads with the alternate allele

DP: Depth (mean coverage across all samples) - aims to eliminate sites with excessive coverage caused by alignment artifacts

Per-sample DP: Depth per individual sample (minimum coverage).

Table S6. Classification results from Discriminant Function Analysis I. Original and predicted group membership results from Discriminant function analysis of *O. korogwe*, *O. niloticus* and identified hybrids in the southern lakes, using traditional methods and geometric morphometric analysis.

|  |  | Classified group | |  |
| --- | --- | --- | --- | --- |
| Measurements | Original group | *O. korogwe* | *O. niloticus* | Total |
| Linear (traditional) | *O. korogwe* | 16 | 2 | 18 |
|  | *O. niloticus* | 1 | 13 | 14 |
|  | Hybrids (OK x ON) | 6 | 2 | 8 |
| Geometric | *O. korogwe* | 17 | 1 | 18 |
| morphometric | *O. niloticus* | 1 | 13 | 14 |
|  | Hybrids (OK x ON) | 4 | 4 | 8 |

Table S7. Correlation of traits with Discriminant Function axes I. Correlation of traits with Discriminant Function Axis 1 separating *O. niloticus* from *O. korogwe* in the southern lakes, using linear (traditional) measurements.

| Trait | Correlation with Axis 1 |
| --- | --- |
| Head Width | 0.533 |
| Head Length | 0.392 |
| Anal fin base length | -0.370 |
| Eye length | 0.367 |
| Body depth | 0.205 |
| Inter orbital width | 0.192 |
| Pelvic fin length | 0.165 |
| Caudal fin length | -0.120 |
| Caudal peduncle length | -0.115 |
| Pectoral fin base length | -0.105 |
| Snout length | -0.080 |
| Dorsal fin base length | -0.058 |
| Caudal peduncle depth | -0.050 |
| Lower Jaw length | -0.013 |

Table S8. Classification results from Discriminant Function Analysis II. Classification results from Discriminant Function Analysis of four populations of *O. korogwe* from a) traditional measures of morphology and b) geometric morphometric measures.

|  |  | Classified group | | | |  |
| --- | --- | --- | --- | --- | --- | --- |
| Measurements | Original group | K-M | K-Z | K-N | K-R | Total |
| Linear (traditional) | O. korogwe Mlingano (K-M) | 31 | 3 | 0 | 0 | 34 |
|  | O. korogwe Zigi (K-Z) | 0 | 23 | 0 | 0 | 23 |
|  | O. korogwe Nambawala (K-N) | 0 | 0 | 13 | 1 | 14 |
|  | O. korogwe Rutamba (K-R) | 1 | 1 | 0 | 7 | 9 |
| Geometric | O. korogwe Mlingano (K-M) | 10 | 0 | 0 | 0 | 10 |
| morphometric | O. korogwe Zigi (K-Z) | 0 | 8 | 1 | 0 | 9 |
|  | O. korogwe Nambawala (K-N) | 0 | 0 | 28 | 1 | 29 |
|  | O. korogwe Rutamba (K-R) | 0 | 0 | 2 | 38 | 40 |

| Trait | Correlation with DF Axis 1 | Correlation with DF Axis 2 | Correlation with DF Axis 3 |
| --- | --- | --- | --- |
| Anal fin base length | 0.148 | -0.036 | -0.045 |
| Body depth | 0.314 | -0.061 | 0.172 |
| Caudal fin length | -0.090 | 0.291 | 0.828 |
| Caudal peduncle depth | 0.446 | -0.049 | 0.157 |
| Caudal peduncle length | 0.089 | 0.133 | -0.105 |
| Dorsal fin base length | 0.169 | -0.050 | -0.276 |
| Eye length | -0.197 | 0.430 | 0.246 |
| Head length | 0.030 | 0.174 | 0.500 |
| Head width | -0.130 | -0.019 | 0.476 |
| Inter-orbital width | 0.226 | 0.458 | 0.337 |
| Lower jaw length | 0.141 | -0.086 | 0.511 |
| Pectoral fin length | 0.365 | 0.031 | 0.149 |
| Pelvic fin length | -0.090 | 0.470 | 0.197 |
| Snout length | 0.016 | 0.338 | 0.318 |

Table S9. Correlation of traits with Discriminant Function axes II. Correlation of traits with Discriminant Function axes separating *O. korogwe* populations, using linear (traditional) measurements.

### Table S10. Data files and number of SNPs by analysis. Data files correspond to code in Supporting Text.

| Description | File | Analysis | SNPs (n) |
| --- | --- | --- | --- |
| Quality-filtered biallelic nuclear SNPs excluding unplaced scaffolds | oreo_nucfiltersnps_biallelic_lg_2perpop.vcf.gz | FST, Dxy, pi | 4,072,183 |
| Quality-filtered biallelic nuclear SNPs excluding *O. placidus* hybrid, SNPs excluding heterozygote only sites | raxml.min4.phy | RAxML ASC | 5,992,590 |
| Quality-filtered biallelic nuclear SNPs LD pruned | oreo_pruned.bim | Admixture | 160,.883 |

**Table S11.** Results of the D3 statistics, testing for statistically significant deviations from expected genetic distance between pairs of individuals.

| **P1** | **P2** | **P3** | **Mean D3** | **SD D3** | **p value** |
| --- | --- | --- | --- | --- | --- |
| Okorogwe__Nambawala__T3J2 | Okorogwe__MlinganoDam__P4A10 | Oniloticus__LakeAlbert__U1A1 | -0.043266683 | 0.002428077 | 0 |
| Okorogwe__Nambawala__T3J2 | Okorogwe__MlinganoDam__P4B1 | Oniloticus__LakeAlbert__U1A1 | -0.031307302 | 0.002423252 | 0 |
| Okorogwe__Nambawala__T3J2 | Okorogwe__MlinganoDam__P4B2 | Oniloticus__LakeAlbert__U1A1 | -0.043255295 | 0.002480584 | 0 |
| Okorogwe__Nambawala__T3J2 | Okorogwe__MlinganoDam__P4A10 | Oniloticus__LakeAlbert__U3A3 | -0.043062768 | 0.002406678 | 0 |
| Okorogwe__Nambawala__T3J2 | Okorogwe__MlinganoDam__P4B1 | Oniloticus__LakeAlbert__U3A3 | -0.031222912 | 0.002447765 | 0 |
| Okorogwe__Nambawala__T3J2 | Okorogwe__MlinganoDam__P4B2 | Oniloticus__LakeAlbert__U3A3 | -0.043104307 | 0.0023835 | 0 |
| Okorogwe__Nambawala__T3J4 | Okorogwe__MlinganoDam__P4A10 | Oniloticus__LakeAlbert__U1A1 | -0.01744169 | 0.001630196 | 0 |
| Okorogwe__Nambawala__T3J4 | Okorogwe__MlinganoDam__P4B1 | Oniloticus__LakeAlbert__U1A1 | -0.005470223 | 0.001669989 | 0.001054365 |
| Okorogwe__Nambawala__T3J4 | Okorogwe__MlinganoDam__P4B2 | Oniloticus__LakeAlbert__U1A1 | -0.017430138 | 0.001641556 | 0 |
| Okorogwe__Nambawala__T3J4 | Okorogwe__MlinganoDam__P4A10 | Oniloticus__LakeAlbert__U3A3 | -0.017495855 | 0.001679343 | 0 |
| Okorogwe__Nambawala__T3J4 | Okorogwe__MlinganoDam__P4B1 | Oniloticus__LakeAlbert__U3A3 | -0.005641 | 0.001544814 | 0.00026064 |
| Okorogwe__Nambawala__T3J4 | Okorogwe__MlinganoDam__P4B2 | Oniloticus__LakeAlbert__U3A3 | -0.017538009 | 0.001620035 | 0 |
| Okorogwe__Nambawala__T3J6 | Okorogwe__MlinganoDam__P4A10 | Oniloticus__LakeAlbert__U1A1 | -0.026655445 | 0.002170778 | 0 |
| Okorogwe__Nambawala__T3J6 | Okorogwe__MlinganoDam__P4B1 | Oniloticus__LakeAlbert__U1A1 | -0.014685469 | 0.002199143 | 2.43E-11 |
| Okorogwe__Nambawala__T3J6 | Okorogwe__MlinganoDam__P4B2 | Oniloticus__LakeAlbert__U1A1 | -0.026642954 | 0.002192584 | 0 |
| Okorogwe__Nambawala__T3J6 | Okorogwe__MlinganoDam__P4A10 | Oniloticus__LakeAlbert__U3A3 | -0.026472375 | 0.002240804 | 0 |
| Okorogwe__Nambawala__T3J6 | Okorogwe__MlinganoDam__P4B1 | Oniloticus__LakeAlbert__U3A3 | -0.014620779 | 0.002181374 | 2.05E-11 |
| Okorogwe__Nambawala__T3J6 | Okorogwe__MlinganoDam__P4B2 | Oniloticus__LakeAlbert__U3A3 | -0.026513275 | 0.002200194 | 0 |
| Okorogwe__Nambawala__T3J2 | Okorogwe__MlinganoDam__P4A10 | Ourolepis__LakeLugongwe__T2J5 | 0.049471943 | 0.00242394 | 0 |
| Okorogwe__Nambawala__T3J2 | Okorogwe__MlinganoDam__P4B1 | Ourolepis__LakeLugongwe__T2J5 | 0.037784552 | 0.002365374 | 0 |
| Okorogwe__Nambawala__T3J2 | Okorogwe__MlinganoDam__P4B2 | Ourolepis__LakeLugongwe__T2J5 | 0.050077397 | 0.00241608 | 0 |
| Okorogwe__Nambawala__T3J2 | Okorogwe__MlinganoDam__P4A10 | Ourolepis__Mbuyunipool__T6A2 | 0.049724793 | 0.00254084 | 0 |
| Okorogwe__Nambawala__T3J2 | Okorogwe__MlinganoDam__P4B1 | Ourolepis__Mbuyunipool__T6A2 | 0.037858938 | 0.002454694 | 0 |
| Okorogwe__Nambawala__T3J2 | Okorogwe__MlinganoDam__P4B2 | Ourolepis__Mbuyunipool__T6A2 | 0.050036855 | 0.002508508 | 0 |
| Okorogwe__Nambawala__T3J4 | Okorogwe__MlinganoDam__P4A10 | Ourolepis__LakeLugongwe__T2J5 | 0.025566503 | 0.001844267 | 0 |
| Okorogwe__Nambawala__T3J4 | Okorogwe__MlinganoDam__P4B1 | Ourolepis__LakeLugongwe__T2J5 | 0.013864769 | 0.001747232 | 2.00E-15 |
| Okorogwe__Nambawala__T3J4 | Okorogwe__MlinganoDam__P4B2 | Ourolepis__LakeLugongwe__T2J5 | 0.02617948 | 0.001897977 | 0 |
| Okorogwe__Nambawala__T3J4 | Okorogwe__MlinganoDam__P4A10 | Ourolepis__Mbuyunipool__T6A2 | 0.025979047 | 0.001885167 | 0 |
| Okorogwe__Nambawala__T3J4 | Okorogwe__MlinganoDam__P4B1 | Ourolepis__Mbuyunipool__T6A2 | 0.014100537 | 0.001754005 | 8.88E-16 |
| Okorogwe__Nambawala__T3J4 | Okorogwe__MlinganoDam__P4B2 | Ourolepis__Mbuyunipool__T6A2 | 0.026296578 | 0.00188004 | 0 |
| Okorogwe__Nambawala__T3J6 | Okorogwe__MlinganoDam__P4A10 | Ourolepis__LakeLugongwe__T2J5 | 0.036884685 | 0.002462671 | 0 |
| Okorogwe__Nambawala__T3J6 | Okorogwe__MlinganoDam__P4B1 | Ourolepis__LakeLugongwe__T2J5 | 0.025187948 | 0.00234499 | 0 |
| Okorogwe__Nambawala__T3J6 | Okorogwe__MlinganoDam__P4B2 | Ourolepis__LakeLugongwe__T2J5 | 0.037491545 | 0.002448547 | 0 |
| Okorogwe__Nambawala__T3J6 | Okorogwe__MlinganoDam__P4A10 | Ourolepis__Mbuyunipool__T6A2 | 0.03715063 | 0.002506768 | 0 |
| Okorogwe__Nambawala__T3J6 | Okorogwe__MlinganoDam__P4B1 | Ourolepis__Mbuyunipool__T6A2 | 0.025275681 | 0.002416399 | 0 |
| Okorogwe__Nambawala__T3J6 | Okorogwe__MlinganoDam__P4B2 | Ourolepis__Mbuyunipool__T6A2 | 0.037463557 | 0.002480429 | 0 |

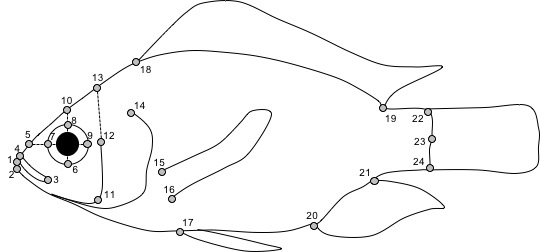

### Figure S1. Landmarks used in the geometric morphometric analysis.

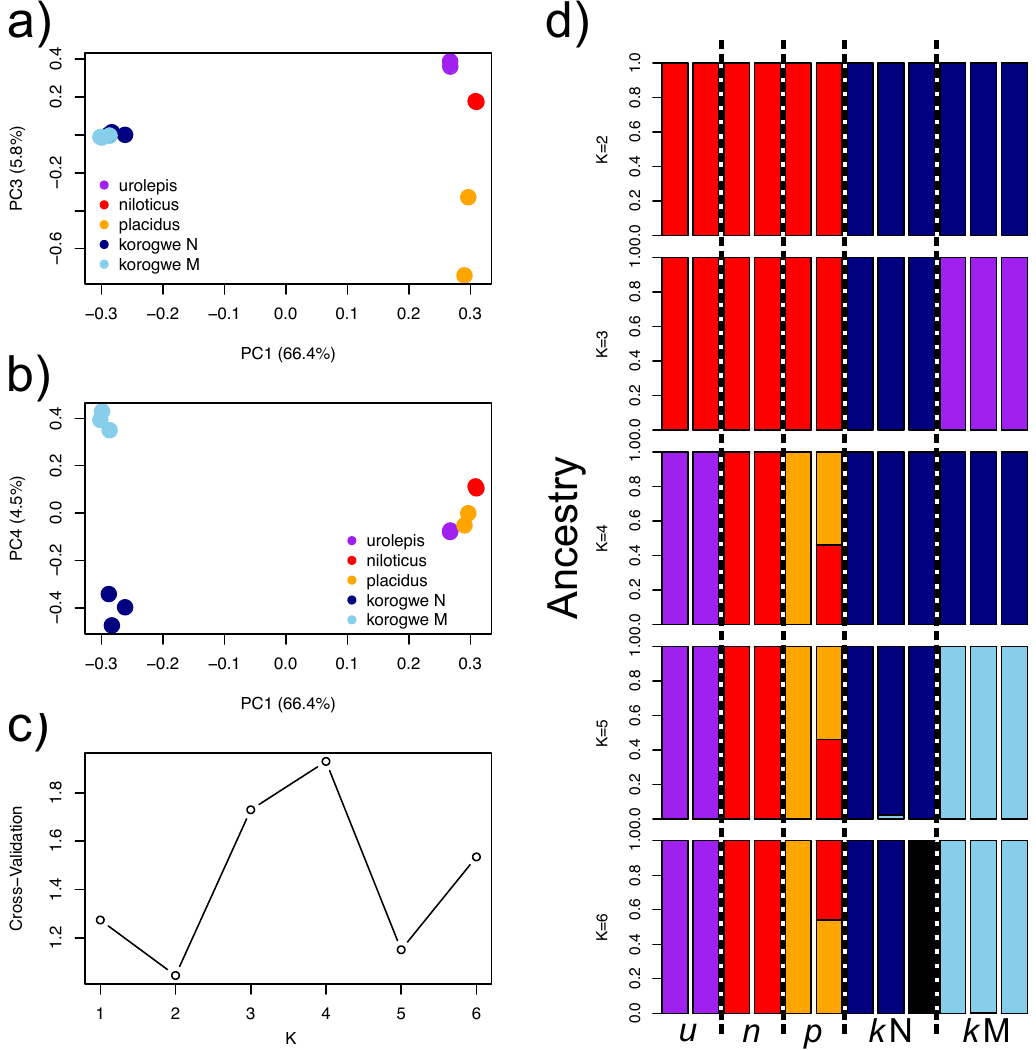

### Figure S2. WGR population genetic and phylogenetic analysis. a-b) PCA analysis of LD-pruned nuclear (116,901) SNPs, c) Cross-validation error for admixture analysis of K=1-6, based on 400,680 LD-pruned nuclear SNPs, d) Admixture cluster membership for K=2-5. Species codes used here are u = *O. urolepis*, n = *O. nilotic*us, p = *O. placidus rovumae*, kN = *O. korogwe* Nambalwala (southern), kM = *O. korogwe* Mlingano northern). Note colours correspond with genetic clusters, and individual colours are selected to best correspond with populations in K=5.

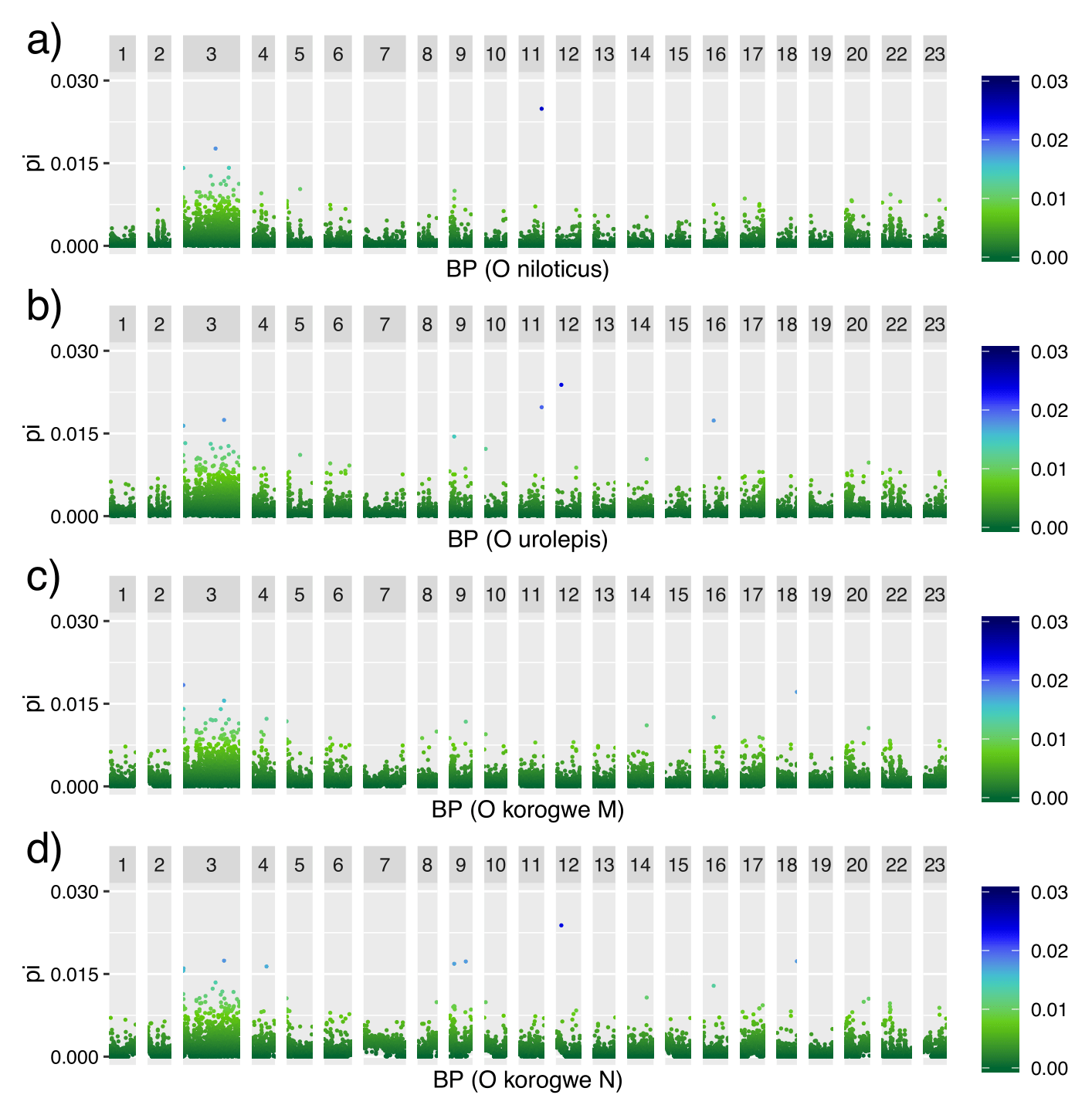

**Figure S3.** Within population nucleotide diversity (pi) across linkage groups, estimated with whole genome data, in non-overlapping 50kb windows.

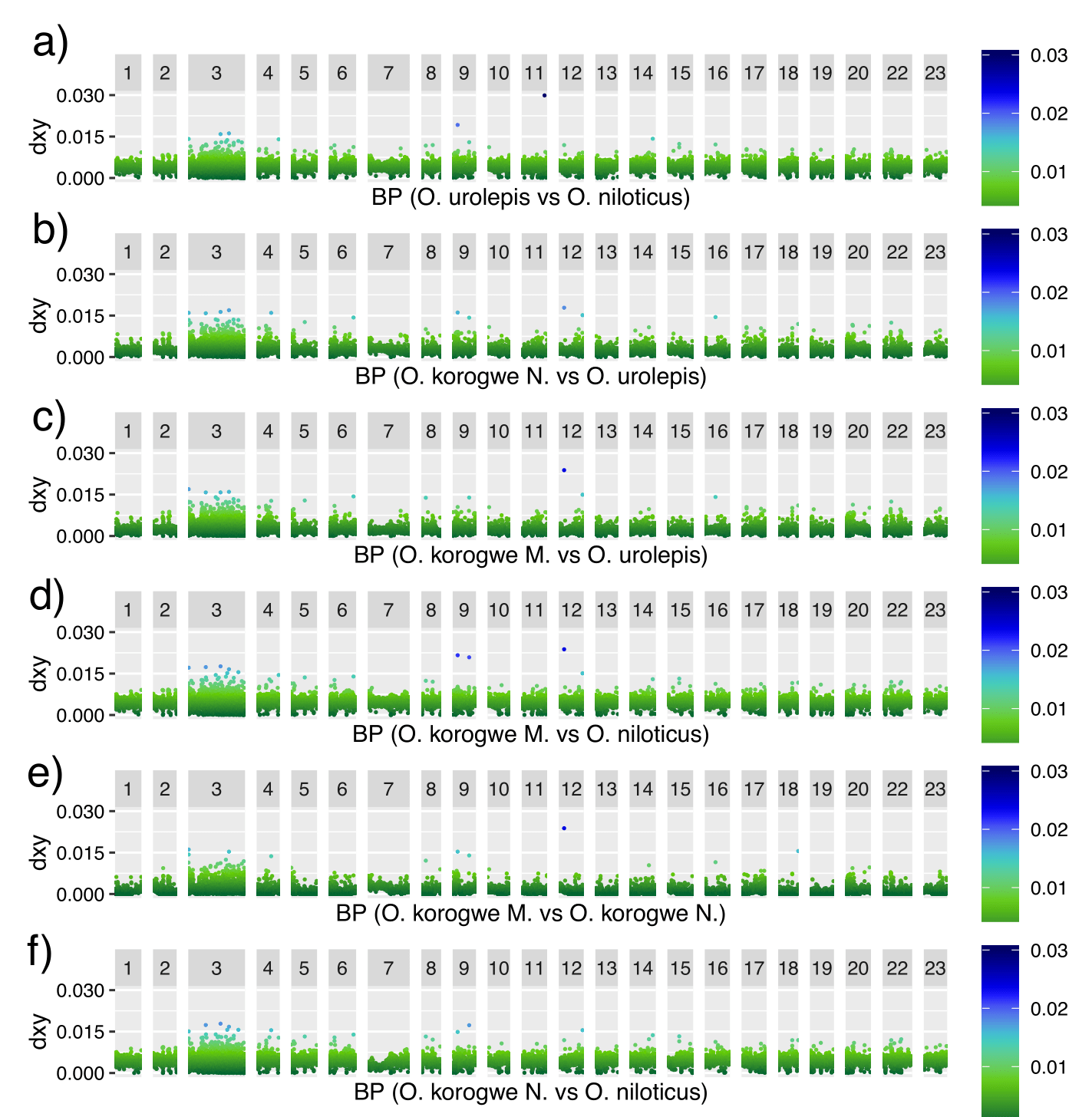

**Figure S4.** Absolute genetic divergence (Dxy) between population pairs across linkage groups, as estimated using whole genome data, in non-overlapping 50kb windows.
